# Controlling malaria mosquito reproduction via the octopamine beta2 receptor

**DOI:** 10.1101/2025.08.20.670845

**Authors:** David A. Ellis, Judy Bagi, Stephen Terry, Eve Doran, Scott Tytheridge, YiFeng YJ Xu, Matthew P. Su, Max Lombardi, Gloria Iacomelli, Matthew Peirce, Melania Ligato, Daniela Terrazas, Watson Ntabaliba, Sarah Moore, Joerg T Albert, Roberta Spaccapelo, Marta Andrés

**Author notes:** Department of Entomology, Cornell University, Ithaca, NY, 14853, USA. The Francis Crick Institute, 1 Midland Road, London, NW1 1AT, UK.

## Abstract

Mosquito reproduction in a broad sense involves multiple steps from acoustic recognition of mating partners to egg hatching. We show that the octopamine receptor AgOctβ2R controls different aspects of this process in a sexually dimorphic manner. *AgOctβ2R* knockout males present auditory defects that impair their ability to inseminate females, whilst knockout females are sterile. These phenotypes suggest AgOctβ2R as a target to impair mosquito reproduction at multiple levels. We test the reproductive effects of the insecticide amitraz, an AgOctβ2R agonist, showing that amitraz exposure reduces insemination in the lab but not in the field and has no effects on female sterility, excluding its applicability as a mating disruptor. Pharmacological assays reveal that AgOctβ2R sensitivity to amitraz is reduced compared to other arthropods, but its responses can be altered by modifying residues in the binding pocket. Together, our results establish AgOctβ2R as a promising target to disrupt mosquito reproduction but emphasize the necessity of developing new tools to exploit this approach.

## Introduction

The spread of insecticide resistance across mosquito populations and their behavioural adaptations to avoid control tools threatens malaria control (*1*). New approaches are needed to identify novel insecticide molecular targets and alternative approaches to fight mosquitoes (*2*). Currently, the most important tools against the malaria mosquito *Anopheles gambiae* are insecticide-treated bed nets and indoor residual spraying. Despite their indisputable success in reducing malaria transmission (*3*), both approaches target only indoor-biting mosquito females, leaving several vector species untargeted. Developing innovative tools to induce behavioural modifications (*4–6*) could complement traditional control methods (*7*, *8*). The mosquito vectorial capacity (its ability to transmit pathogens) relies on multiple parameters, including female mosquito population size, which is largely dependent on reproductive success. Blocking mosquito reproduction is thus an alternative approach for mosquito control that is increasingly being explored (*9–11*).

Mating of *An. gambiae* takes places in swarms (*12*, *13*), where males acoustically detect females through their flight tones before chasing them to copulate (*14–17*). Interrupting the precopulatory acoustic recognition has been proposed as an innovative target for mosquito control (*18*, *19*). The highly complex physiology of mosquito audition offers multiple targets for manipulating mosquito hearing (*20*). Genetically disrupting mosquito audition has been recently shown to impair mating in the yellow fever mosquito *Aedes aegypti* (*21*). However, molecular pathways suitable for a pharmacological disruption of mosquito hearing to develop mating disruptors have not yet been identified.

We recently showed that octopamine modulates the auditory mechanics of *An. gambiae* through activation of the β2-adrenergic-like octopamine receptor, AgOctβ2 (*22*). As octopamine receptors are found exclusively in arthropods (*23*), they are promising targets for novel insecticides (*24–26*). Currently, the formamidine amitraz is the only available insecticide that targets octopamine receptors (IRAC group 19) (*27*). Amitraz is widely used as a pesticide in agriculture against ticks and parasitic mites (*28*, *29*). *In vivo*, amitraz targets β2-adrenergic octopamine receptors, although sensitivity varies across arthropods (*30*). This differential sensitivity of amitraz has been exploited to selectively target pests without affecting non-target pollinators (*30*). The potential for using amitraz to control mosquito populations has not been thoroughly explored, possibly due to its low toxicity (*31*, *32*). In *An. gambiae*, amitraz activates AgOctβ2 and induces the erection of antennal fibrillae (*22*), suggesting that targeting AgOctβ2 could be exploited for mosquito reproductive control.

In this paper, we show that the auditory defects of *AgOctβ2* knockout male mosquitoes cause severe mating behaviour defects. Moreover, via an auditory-independent mechanism, *AgOctβ2* knockout females are sterile. This establishes AgOctβ2 as a promising candidate to target different aspects of mosquito reproduction: male mate-seeking and female fertilisation. We also assessed potential reproductive effects of amitraz, showing that it only causes a mild reduction of female insemination rates, but has no effects on female fertility. Additionally, we performed *in vitro* studies to compare Octβ2 receptor responses to amitraz across arthropods, demonstrating that the *An. gambiae* receptor is less sensitive than other susceptible orthologs. Altogether, our results provide strong evidence to promote AgOctβ2 as a candidate for malaria mosquito reproductive control but also highlight the necessity to develop novel tools for vector control applications.

## Results

### Octopamine modulates the auditory tuning and sensitivity of *An. gambiae* through AgOctβ2 signalling

The mosquito ear is composed of the flagellum, which acts as sound receiver, and the auditory organ, or Johnston’s organ (JO), located at the flagellar base (*12*) (Fig. 1a). Flagellar vibrations caused by sound emissions are transduced into electrical signals by auditory neurons in the JO. Interestingly, the mechanical best frequency (the stimulus frequency which induces the largest flagellar mechanical vibrations) and the electrical best frequency (the stimulus frequency which induces the largest nerve responses) do not match in male *An. gambiae*, reflecting its distortion product-based hearing (*15*, *17*, *33*). We recently showed that octopamine modulates mechanical aspects of male mosquito audition, increasing the flagellar stiffness and tuning frequency (*22*). To gain a more holistic understanding of octopamine in male audition, we analysed its effects on the electrical activity of the ear.

**Figure 1:**
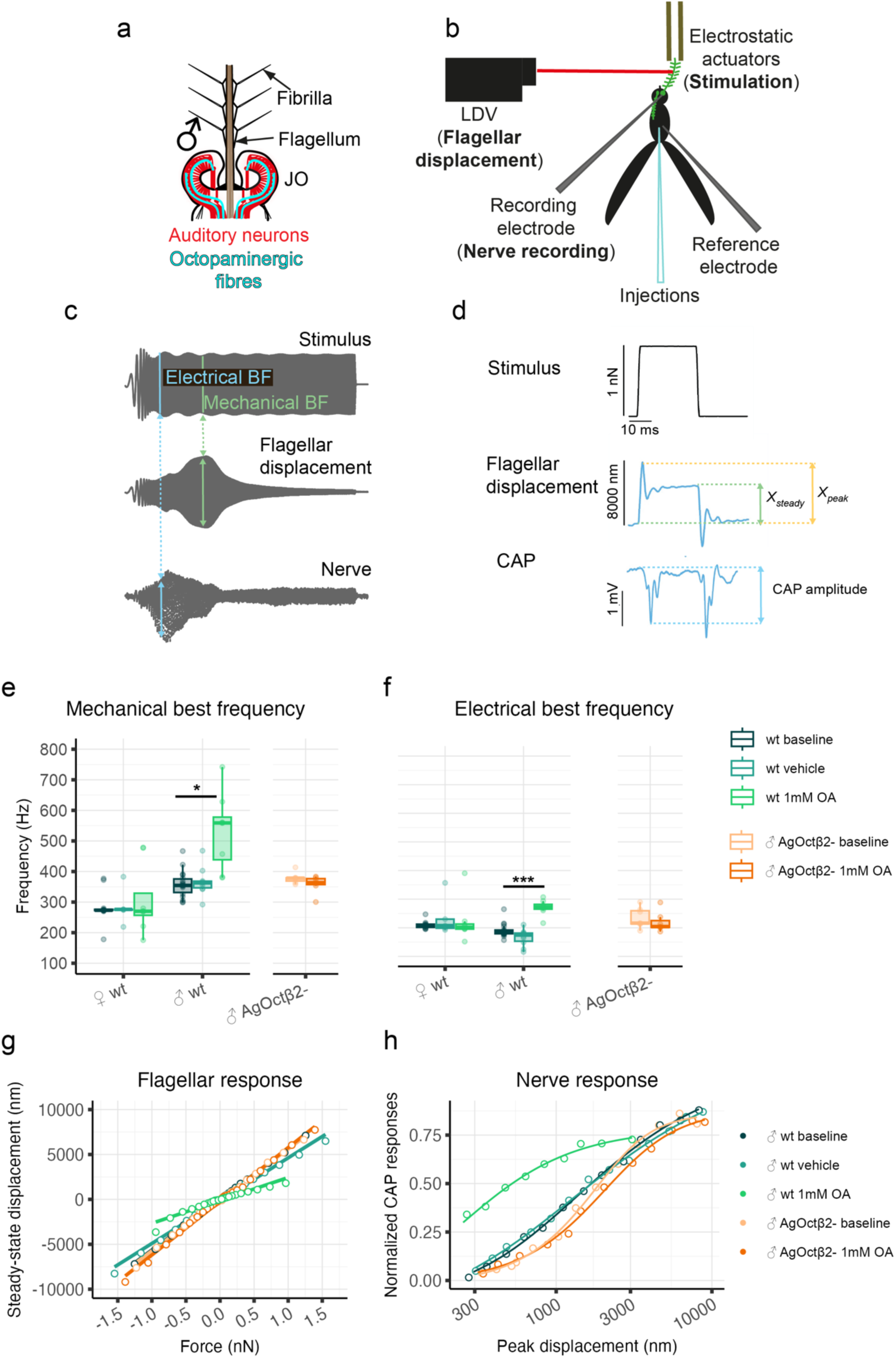
Disrup.ng AgOctβ2 signalling causes auditory defects in male malaria mosquitoes. **a**) Schematic of the malaria mosquito ear (left). Air particle vibrations caused by a sound wave deflect the flagellum, which acts as sound received. At the base of the flagellum, the auditory organ or Johnston’s organ (JO), houses the auditory neurons that transduce the mechanical stimuli into electrical signals. The JO is innervated by octopaminergic neurons that travel from the brain and release octopamine. **b**) Schematic of the experimental setup for auditory tests (right). The laser beam of a laser doppler vibrometer (LDV) is pointed at the tip of the right flagellum to record flagellar displacement. A recording electrode inserted in the antennal nerve records nerve activity.

We used a laser doppler vibrometer (LDV) to record flagellar vibrations (displacements), and an electrode inserted in the antennal nerve to measure compound action potential (CAP) responses (Fig. 1 b). We provided electrostatic actuation to mechanically stimulate the flagellum (Fig. 1c, frequency-modulated sweeps, 0-1000 Hz in 1s). We extracted the mechanical and electrical best frequencies, and compared them before and after octopamine injection in wild-type (wt) male and female mosquitoes (Fig. 1e). We confirmed an increase in the mechanical tuning frequency upon octopamine injections in males but not females (Fig. 1e and Suppl. Table 1, female_baseline_: 282 ± 49 Hz showing mean ± standard deviation, female_octopamine_: 305 ± 112 Hz, Wilcoxon rank-sum test: p > 0.05; male_baseline_: 358 ± 43 Hz, male_octopamine_: 533 ± 124 Hz, t-test p < 0.05). In addition, antennal nerve electrophysiological recordings showed that the electrical tuning also shifted to higher frequencies after octopamine injection, but again only in males (Fig. 1f, female_baseline_: 282 ± 49 Hz, female_octopamine_: 221 ± 71 Hz, Wilcoxon rank-sum test: p > 0.05; male_baseline_: 166 ± 29 Hz, male_octopamine_: 271 ± 27 Hz, Wilcoxon rank-sum test p < 0.01). These results show that the effects of octopamine on male mosquito hearing manifest as electrical changes, and therefore impact sound perception.

We repeated these experiments in *AgOctβ2* knockout male mosquitoes(*22*). Knockout males did not show any change in auditory tuning upon octopamine injection (Fig. 1e,f and Suppl. Table 1). The mechanical tuning remained at ∼350 Hz (*AgOctβ2^-^_baseline_*: 378 ± 18 Hz, *AgOctβ2^-^ _octopamine_*: 359 ± 28 Hz, Wilcoxon rank-sum test: p > 0.05), whilst the electrical tuning remained at ∼250 Hz (*AgOctβ2^-^_baseline_*: 235 ± 36 Hz, *AgOctβ2^-^_octopamine_*: 219 ± 34 Hz, t-test p>0.05). We concluded that octopamine mechanical and electrical auditory effects are mediated by AgOctβ2.

Next, we analysed potential roles of octopamine in modulating the auditory sensitivity. We applied force-step stimulation of decreasing amplitude to the mosquito flagellum and measured resulting flagellar and nerve responses. Step-actuation induces a characteristic response in the flagellum, including an initial overshoot in the positive direction (peak displacement) followed by a rebound and a subsequent positive displacement until a steady-state displacement is reached (steady-state displacement, Fig. 1d). We studied the relationship between the strength of the electrostatic force applied to the flagellum tip and the resulting steady-state flagellar displacements. Octopamine injection in wt individuals caused a decrease in displacement by force, indicating an increased steady-state stiffness (*22*) that was not shown by *AgOctβ2^-^* individuals (Fig. 1g). We also analysed octopamine effects on the amplitude of electrical responses (Fig. 1h, Supp. Fig. 1). Plotting median normalised CAP responses ((V- V_min_)/(V_max_-V_min_)) against flagellar displacements showed a strong enhancement of CAP amplitudes upon octopamine injection, suggesting that, independently of its effects on the antennal mechanics, octopamine enhances the electrical signals generated by auditory neurons. This effect was not observed in *AgOctβ2* knockout males.

Electrostatic actuation is used to provide mechanical stimulation. **c**) Example of frequency-modulated sweep stimulation that was applied to the mosquito flagellum and corresponding flagellar and nerve responses. The stimulus (top) consisted of a linear frequency sweep that ranges from 0 Hz to 1. Maximum amplitude flagellar and nerve responses were identified and corresponding stimulus frequency that elicited those largest responses was extracted as mechanical and electrical best frequency. **d**) Example of force-step stimulation and corresponding flagellar and nerve responses. Force applied to the flagellum causes an initial overshoot in the direction of the force (peak displacement), followed by a rebound and a damped oscillation until a steady-state displacement is reached (*X_steady_*). In the nerve, CAPs are generated with stimulus onset and offset. **e**) Box plots representing the mechanical best frequencies. In wt mosquitoes (green), octopamine injections caused a shift of mechanical best frequency (from ∼ 350 Hz to ∼ 530 Hz, t-test p-value < 0.05), but only in males. Knocking out *AgOctβ2* (orange) abolishes the tuning frequency increase (Wilcoxon rank-sum test, p-value > 0.05), compared to wt mosquitoes. Box plots represent: central line, median; box limits, first and third quartiles; lower and upper whiskers, 5th and 95th percentiles, respectively. **f**) Box plots representing the electrical best frequencies. Octopamine injections cause an increase of the electrical best frequency in male malaria mosquitoes (green, from ∼ 160 Hz to ∼ 270 Hz, Wilcoxon rank-sum test, p-value <0.01), which is not observed in *AgOctβ^-^* individuals (orange, Wilcoxon rank-sum test, p-value > 0.05). Note that mechanical and electrical best frequencies differ in malaria mosquitoes(*17*, *33*). **g**) Steady-state displacement as a function of the external force (linear fit). Octopamine injections in wt individuals cause an increase in the steady-state stiffness (*22*), as the displacement caused by force-steps of same magnitude decreases. Vehicle injections or octopamine injections in *AgOctβ2^-^* males do not cause changes in the amplitude of mechanical responses. **h**) Median antennal nerve normalized CAP amplitudes ((V-V_max_)/(V_max_-V_min_)) in response to flagellar displacements (5 parameter log-logistic fits). A strong increase in CAP amplitude relative to flagellar displacement values is observed upon octopamine injections in wt individuals compared to all other conditions. Samples sizes= 8. BF best frequency, CAP compound action potential, LDV Laser Doppler vibrometer, OA octopamine, wt wild-type.

### Knocking out AgOctβ2 in *An. gambiae* mosquito males causes defects in mating behaviour

Our next question was if the auditory phenotypes of *AgOctβ2* mutants caused behavioural effects. Male *An. gambiae* are attracted to sounds mimicking female wingbeats (phonotactic behaviour). To test effects on phonotaxis, we exposed male mosquitoes to pure tones of different frequencies (200-750 Hz, female wingbeat frequency ∼500Hz) during swarm time. We observed reduced attraction rates of *AgOctβ2^-^* males compared to wt individuals (Fig. 2a, GLM (binomial), p<0.001). These results show that impairing AgOctβ2 signalling affects auditory precopulatory behaviours.

**Figure 2:**
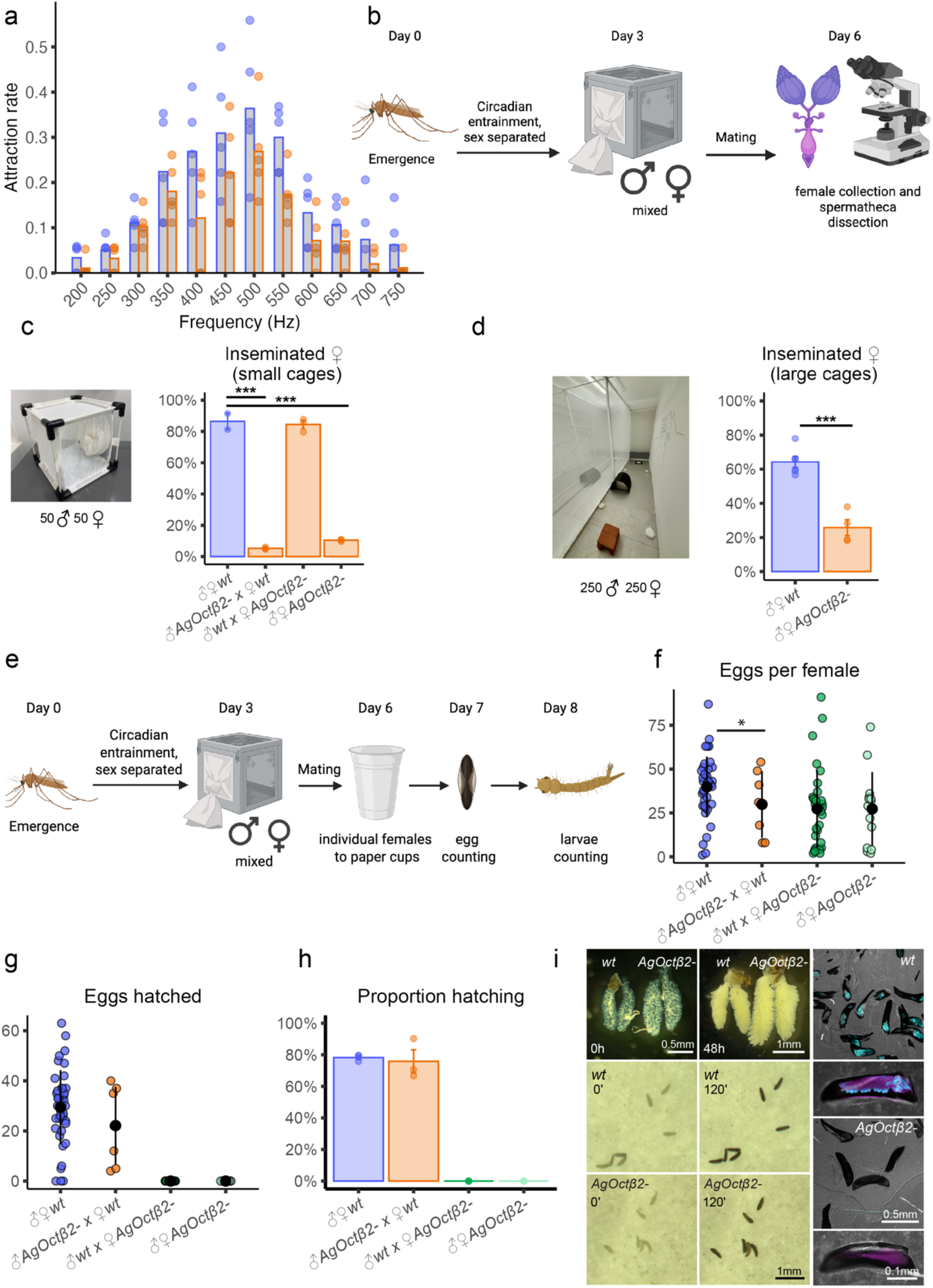
Disrup.ng AgOctβ2 signalling in *An. gambiae* severely affects mating behaviour and causes female sterility. **a**) Phonotactic (attraction) responses of males to biologically relevant sound frequencies (200-750 Hz). *AgOctβ2^-^* males show reduced attraction rates compared to wt mosquitoes (GLM (binomial), p<0.001). Each dot represents the mean of a replicate using 20 mosquitoes. **b**) Experimental design for mating behaviour assays where mosquitoes were allowed to mate for two swarming periods created using BioRender.com. **c**) Crosses in small cages involving *AgOctβ2^-^* males showed reduced female insemination rates, independently of the female genotype (GLM (binomial), p<0.001). *AgOctβ2* knockout in females did not affect mating behaviour. Each dot represents the mean of a replicate using 50 mosquitoes, total of 3 replicates. **d**) Likewise, experiments in large cages (right) also showed a reduction of mating rates when male and female AgOctβ2-mosquitoes were compared to wt crosses. Each dot represents the mean of a replicate from 100 dissected females. **e)** Experimental design for female fertility assays. Females were placed in individual cups after 72h of mating. Numbers of eggs laid, and larvae were counted. **f**) The number of eggs per female were reduced in crosses where *AgOctβ2^-^* males were crossed to wt females (GLM (binomial), p<0.05), but not differences were observed for other genotypes combinations (GLM (binomial), p>0.05). **g**) Plots showing the number of hatched eggs (L1) for each cross. No larvae were found in cups containing *AgOctβ2^-^* females, indicating female sterility, also represented as hatching rates (**h**), where each dot represents the mean of a replicate using 50 females in the initial crosses, total of 3 replicates**. i)** Ovary and egg development in *wt* and *AgOctβ2^-^* mutant females. Top: Ovaries before blood meal (left) and 48 hours after (right); middle: wt eggs immediately (left) and 120 minutes (right) after laying; bottom: *AgOctβ2^-^* mutant eggs immediately (left) and 120 minutes (right) after laying. (Right) Eggs stained with DAPI (nuclear marker); wt eggs present positive DAPI staining, indicating cell division characteristic of the developmental stage. By contrast, *AgOctβ2^-^* eggs are not labelled by DAPI, suggesting that they have not been fertilised.

As phonotactic responses reflect the ability of males to localize females during courtship behaviour (*33*), we next tested the impact of knocking out *AgOctβ2* on mating (Fig 2b). We conducted mating experiments in small cages (30 cm x 30 cm) by mixing 50 males and 50 females of wt or *AgOctβ2^-^* genotypes in different combinations. We observed that the mating rates of crosses with *AgOctβ2^-^* males were severely reduced compared to crosses in which males were wt (Fig. 2c; ♂♀ *wt*: 86 ± 7.2%, ♂ *AgOctβ2^-^* x ♀ *wt*: 5% ± 0.7%, ♂ *wt* x ♀ *AgOctβ2^-^*: 84 ± 4.3 %, ♂♀ *AgOctβ2^-^*: 10 ± 0.7 %, GLM (binomial), p<0.001). This effect was independent of the female genotype. We repeated the experiments in large cages where mosquitoes can form larger swarms. In this case, we compared the effects of crossing *wt* or *AgOctβ2*- males and females. The reduced mating rate of knocked out *AgOctβ2* individuals was also observed in these conditions (Fig.2d; ♂♀ *wt*: 64.2%, ♂♀ *AgOctβ2-*: 26.0%, GLM (binomial), p<0.001), although the phenotype was milder than in smaller cages.

### AgOctβ2 is essential for female fertility

As octopamine is implicated in many aspects of female reproduction across insects (*34–37*), we examined phenotypes related to egg production in *AgOctβ2^-^* females (Fig. 2e). We crossed 50 males and 50 females of different genotypes and counted the number of eggs laid per female and their hatching rate. Crosses of *AgOctβ2^-^* males and wt females showed a reduced number of eggs laid per female compared to control crosses (Fig. 2f; ♂♀ *wt*: 39.5 ± 3.47, ♂ *AgOctβ2^-^*x ♀ *wt*: 29.7 ± 6.6, GLM (binomial), p<0.05), but there were no differences in other crosses (Fig. 2f; ♂ *wt* x ♀ *AgOctβ2^-^*: 26.7 ± 7.4, ♂♀ *AgOctβ2^-^*: 28.9 ± 2.1, GLM (binomial), p>0.05). However, none of the eggs oviposited by mutant females hatched, regardless of the male genotype (Fig. 2g,h; ♂♀ *wt*: 76.6 ± 1.8 %, ♂ *AgOctβ2^-^*x ♀ *wt*: 75.8 ± 7.3 %, ♂ *wt* x ♀ *AgOctβ2^-^*: 0%, ♂♀ *AgOctβ2^-^*: 0%). Phenotypical characterisation of embryos (Fig.2i) showed a lack of DAPI (nuclear) staining in eggs laid by *AgOctβ2^-^* females eggs, suggesting lack of fertilisation. Consequently, *AgOctβ2^-^* females are sterile.

### A novel *in-vivo* screening platform for AgOctβ2 antagonists: current compounds show no effects on mosquito insemination rates

We then examined potential approaches to disrupt AgOctβ2 signalling pharmacologically using feeding assays that can be implemented in behavioural tests. To identify compounds that block AgOctβ2, we exploited the simple physiological process of fibrillae erection in *An. gambiae* males, which is under AgOctβ2 control (*22*). We fabricated 24-well plates where mosquitoes could be monitored with a camera from above as they fed in the dark (Fig. 3a). The proportion of video frames (percent of time) where a given mosquito had erect fibrillae was used as a proxy for AgOctβ2 activation (Fig. 3b). As a proof of concept, we fed a range of octopamine concentrations to mosquitoes and manually categorised them as having erect, partial, or collapsed fibrillae (Fig. 3c). We found a clear dose-response in the percent of time fibrillae were erect (or partially erect) (Fig. 3d).

**Fig 3:**
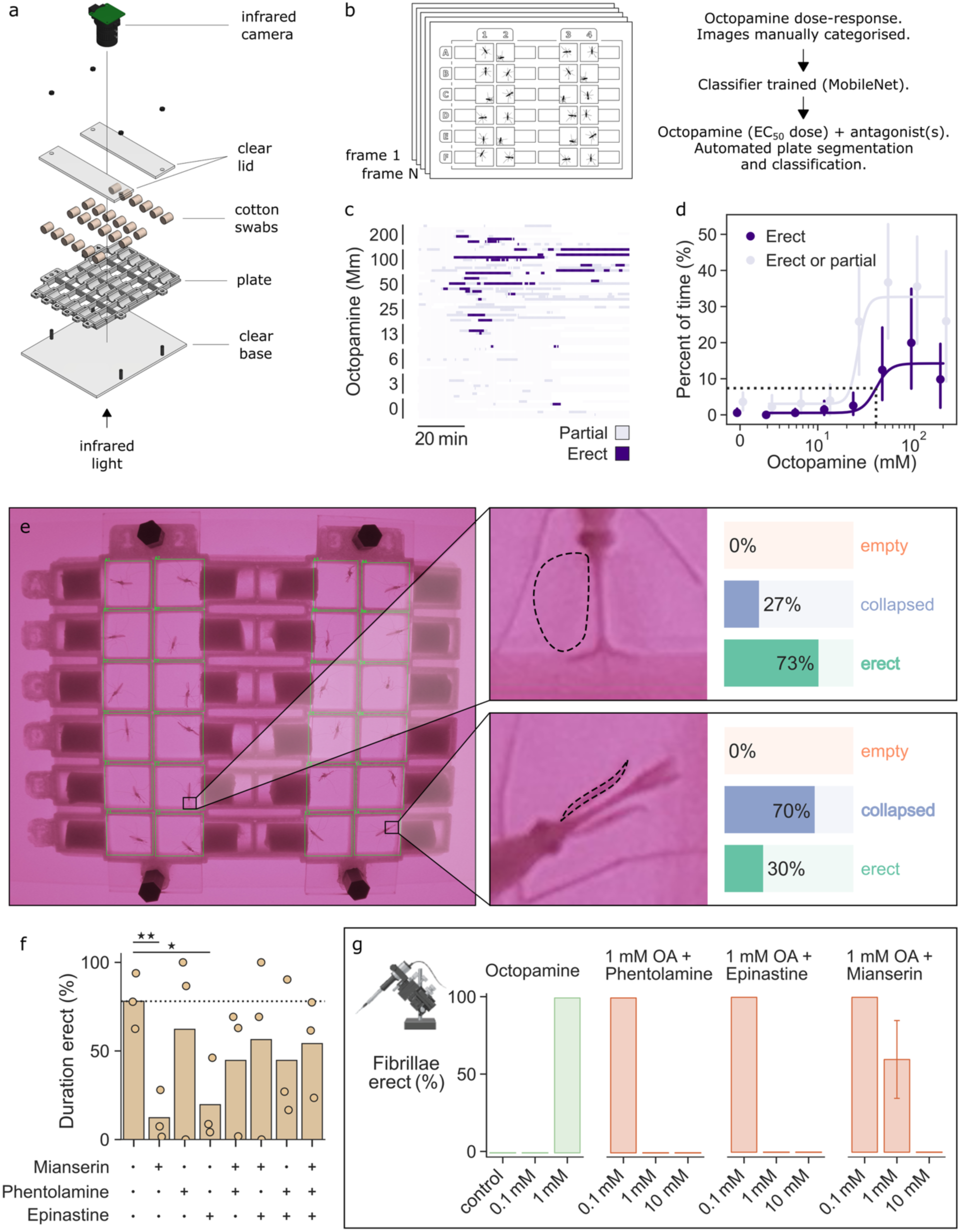
Feeding assay to test the effects of different beta-adrenergic-like octopamine receptor antagonists on fibrillae erection inhibition. **a**) Explosion diagram of the 24-well plates fabricated for this study. Small volumes of drug in sugar solution were provided to mosquitoes via the soaked cotton swabs to the side of each well. Wells were transparent on both top and bottom. To prevent condensation obscuring images, lids were made to cover just the wells and not the swabs. **b**) After an experiment, the status of each mosquito’s fibrillae can be categorised in each frame. This was performed manually on an initial experiment with varying concentrations of octopamine, with the resulting data providing both an EC50 dose for antagonist feeding experiments, as well as training data for an image classifier. This classifier, along with a pipeline to automatically segment plate images into individual wells, was then used to analyse experiments testing the efficacy of various Octb2 antagonists. **c**) Raster plot showing the fibrillar dynamics of several mosquitoes over the course of an experiment. Octopamine was provided to the mosquitoes at a range of doses, and the status of fibrillae was categorised manually. Deep purple represents video frames where a mosquito showed clear fibrillae erection. Light purple frames show frames where fibrillae were deemed to be partially erect. Each row represents a single mosquito and the x-axis represents time. **d**) Dose-response profile of fibrillae at a range of octopamine concentrations. Each point represents the average percentage of frames those mosquitoes showed erect (deep purple) or erect and partially erect (light purple) fibrillae. Raw data is shown in c. The EC50 concentration was determined using only erect data (purple) and is shown by the dotted lines. Four parameter logistic regressions were fitted to all data bar the negative control (far left). **e**) Example of a video frame where wells have been segmented (green boxes) and an image classifier was run on each sub-image to determine fibrillae status. Two mosquitoes have been highlighted: one where the classifier predicted fibrillae were erect, and one where it predicted they were collapsed. The image classifier (MobileNet CNN, Google Teachable Machine) was trained on ∼750 images per class, including augmentations. **f**) Predicted fibrillae status of mosquitoes fed three Octb2 antagonists, on their own or in combination. Antagonists were provided in sugar alongside the EC50 dose of octopamine inferred in d (36.7 mM). Drug concentrations were based on their maximum solubility (mianserin: 5 mM; phentolamine: 20 mM; Epinastine: 10 mM). Fibrillae status was predicted using the pipeline described in e. Each point represents a mosquito; bars show the mean. **g**) Injection assays to validate feeding test. Different octopamine concentrations were injected or co-injected three Octβ2 receptor antagonists (phentolamine, epinastine and mianserin). Equimolar concentrations of phentolamine and epinastine counteract 1mM octopamine effects. Three replicates were conducted, 5 mosquitoes were tested in each replicate.

The manually curated octopamine data also had two further benefits. Firstly, an EC_50_ dose, where the fibrillar response was 50% of maximum, was interpolated from the more conservative erect-only dataset (Fig. 3d). This dose (36.7 mM) provided a useful point at which to test the efficacy of antagonists at blocking octopamine-dependent fibrillae erection. Secondly, the manually categorised images provided a data set that could be used to train an image classifier. As well as images of empty wells (to account for background), this model was trained on erect and collapsed images. The accuracy of each class was as follows: empty=0.99 (100 test images), erect=0.81 (114 test images), collapsed=0.78 (115 test images). This model was then integrated into a pipeline where plate images were segmented into wells, before being classified as erect, collapsed or empty (Fig. 3e). We tested the efficacy of three known antagonists (*38*) at preventing fibrillae erection when co-fed with the EC50 dose of octopamine, on their own and in combination. As this was an exploratory approach, we used high doses of antagonist, basing the concentrations on how soluble they each were (mianserin: 5 mM; phentolamine: 20 mM; Epinastine: 10 mM). Mianserin and epinastine on their own, but not in any combination, significantly blocked fibrillae erection when compared to the no-antagonist control (independent two-sample t-tests: mianserin, T=5.432, p=0.006; epinastine, T=3.627, p=0.022).

To corroborate the results of our *in vivo* screening platform, we administered the same compounds using thoracic injections. Firstly, we observed that 1 mM octopamine caused fibrillae erection in 100% of mosquitoes. Note that although it is expected that compound injection shows stronger effects than feeding, it can damage mosquitoes hindering behavioural applications. We co-injected 1 mM octopamine with different antagonist concentrations (Fig. 3g). We observed that, in line with feeding assay results, all three antagonists counteracted octopamine effects, but in this case epinastine and phentolamine were more potent. We also test an alternative administration method by topically applying the compounds dissolved in 80% DMSO on the mosquito thorax. Topical assays required 100 mM octopamine to induce fibrillae expansion in 100% of the mosquitoes (Suppl. Fig. 2), demonstrating lower efficacy as a delivery method compared to compound feeding or injection.

Following the feeding assay results, we selected mianserin and epinastine to assess their effects on mosquito mating behaviour (Fig. 4a). Fifty male and 50 female mosquitoes were starved, mixed and fed on either 5 mM mianserin or 2.5mM epinastine for 22 hours (including one swarming period). Females were collected and their insemination status assessed. We did not observe significant differences between treatment and control groups for either antagonist (Fig. 4b, insemination rate, control_epinastine_: 47.3 ± 10.2%, 2.5 mM epinastine: 37.7 ± 3.9%; control_mianserin_: 52.1 ± 3.2%, 5 mM mianserin: 58.8 ± 5.6%). However, we observed high mortality across males in the treatment cage for both drugs (Suppl. Fig. 3; control_epinastine_ female: 7 ± 3%, 2.5 mM epinastine female: 5 ± 0.05 %, control_epinastine_ male: 16 ± 4 %, 2.5 mM epinastine male: 46 ± 2 %; control_mianserin_ female: 8.0 ± 2.0%, 5 mM mianserin female: 8.0 ± 2.0 %, control_mianserin_ male: 3.0 ± 1.0 %, 5 mM mianserin male: 24.5 ± 0.5 %). Due to the high mortality induced by both drugs, we stopped analysing their effects on mating behaviour as results were difficult to interpret.

**Figure 4:**
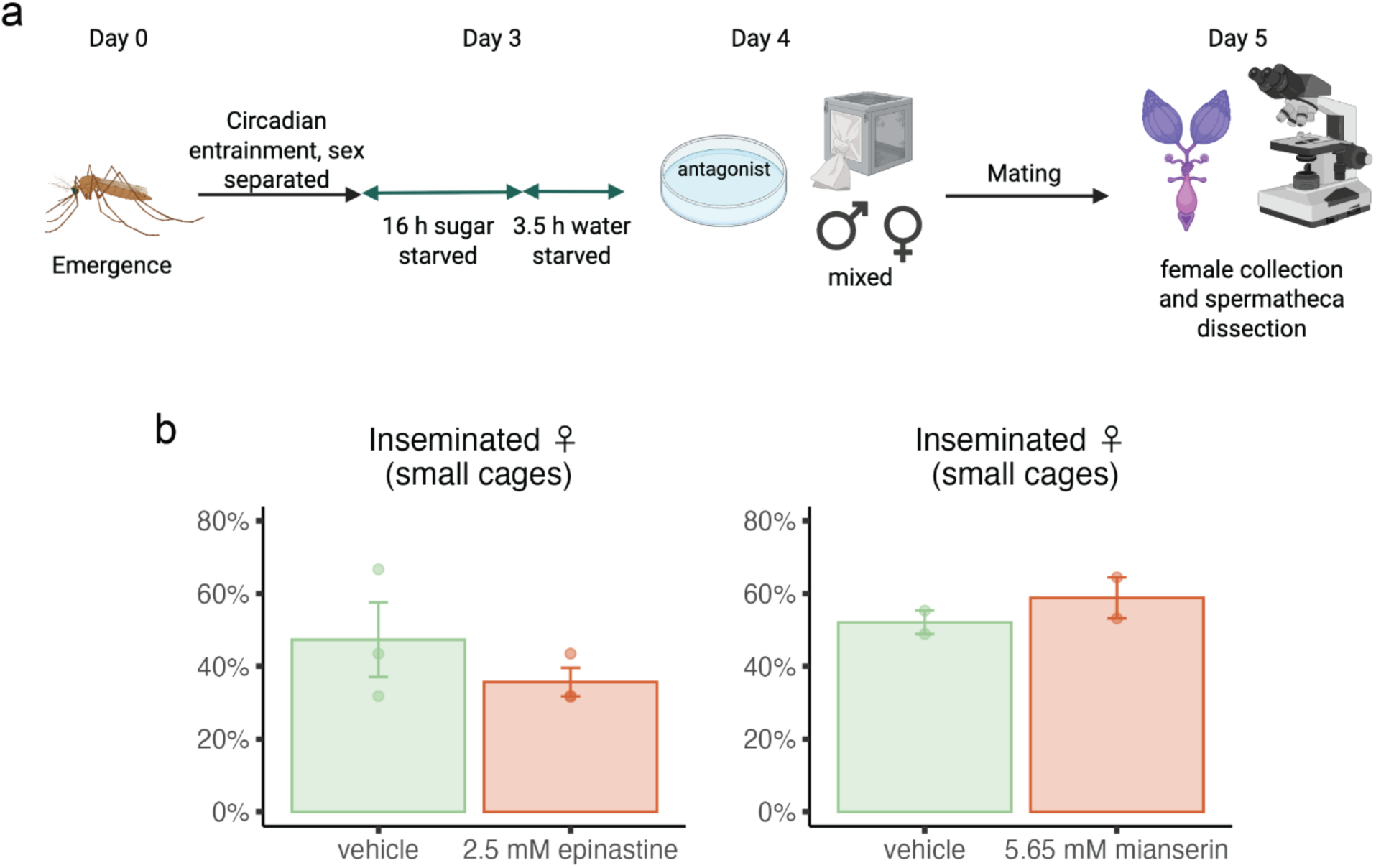
Feeding mosquitoes with beta-adrenergic-like octopamine receptor antagonists do not affect mating behaviour. **a)** Experimental design for antagonist feeding mating assays created using BioRender.com. **b**) Feeding mosquitoes with neither 2.5mM epinastine or 5 mM mianserin did not affect female insemination rates compared to vehicle feeding (GLM (binomial), p>0.05). Each dot represents the mean of a replicate using 50 females in the initial crosses, total of 3 (epinastine) and 2 (mianserin) replicates.

### Targeting AgOctβ2 using amitraz subtly affects mosquito insemination rates

We then decided to test the behavioural effects of the insecticide amitraz, which acts as an *in vivo* agonist of AgOctβ2, and causes fibrillae erection in wild-type, but not *AgOctβ2^-^,* males (*22*). Here we further tested long-term effects of amitraz on fibrillae erection, along with locomotion and mortality in *An. gambiae*. We exposed 25 male mosquitoes to three different concentrations of amitraz (0.1%, 0.4%, 4%) for 3 min, consistent with WHO cone bioassays (*39*).

All three amitraz concentrations caused fibrillae erection in almost 100% of male mosquitoes (Fig. 5a), and this effect lasted for one hour in more than 70% of males (control: 0%, 0.1% amitraz: 78.1 ± 20.2%, 0.4% amitraz: 68.5 ± 28.0%, 4% amitraz: 95.7 ± 9.4%). Even after 24 h, around 20% showed erect fibrillae. We also tested potential knockdown effects of amitraz (Fig. 5b), observing negligible effects on mosquito mobility after one hour (mosquito knockdown: control: 0%, 0.1% amitraz: 1.0 ± 4.4%, 0.4% amitraz: 1.8 ± 5.7%, 4% amitraz: 0%), validating the use of amitraz for further mating behavioural tests. We also tested effects on mosquito survival, as we could not find any such study conducted in *An. gambiae*. We observed a similar survival distribution in control and 0.4% amitraz treated mosquitoes (Fig. 5c), confirming that amitraz is not a suitable insecticide to kill mosquitoes.

**Figure 5:**
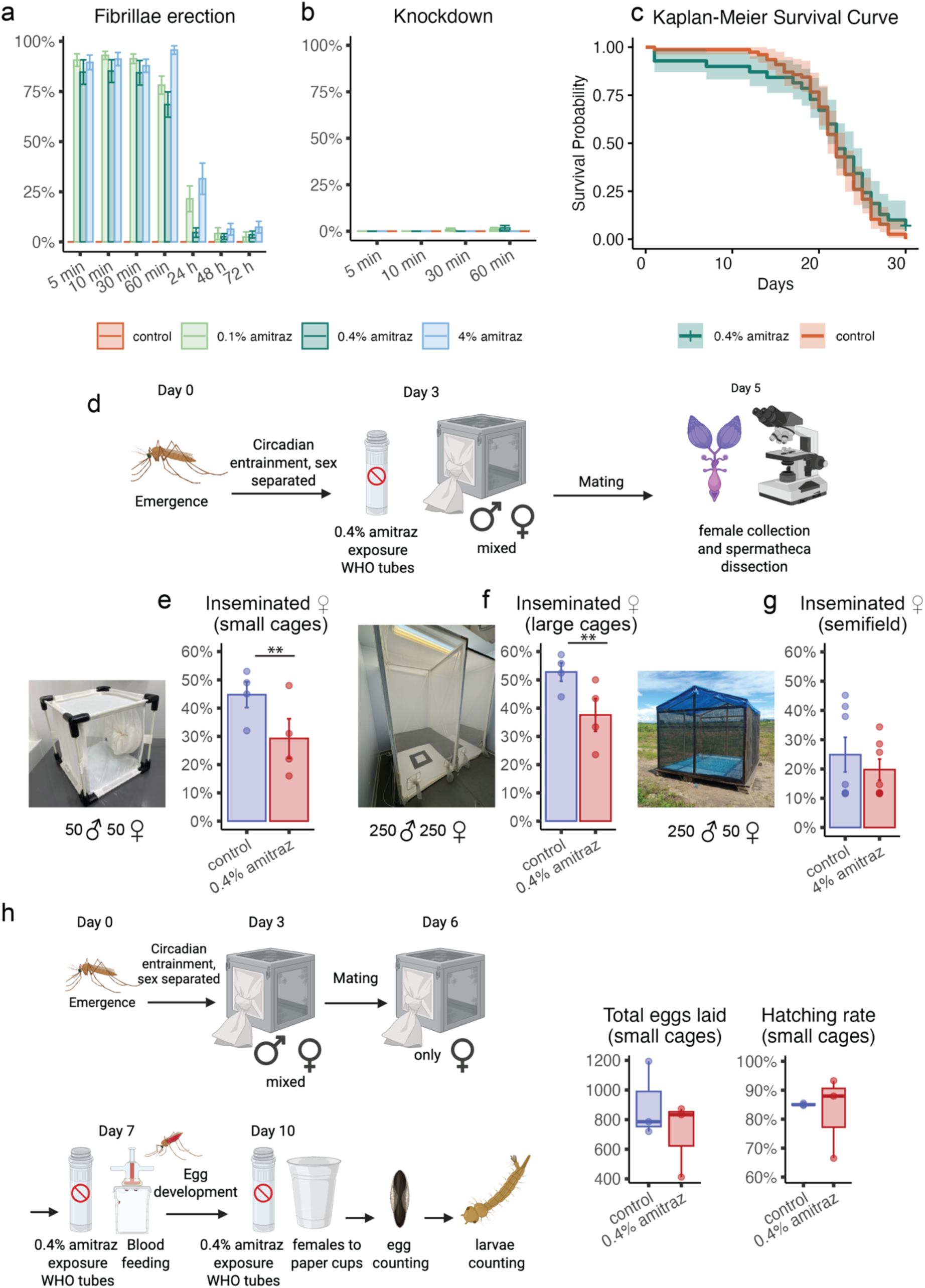
Amitraz exposure reduces female insemination rates in the lab, but has no effects in the field or in female sterility. **a)** Effects of amitraz exposure on the erection of antennal fibrillae in male mosquitoes. The three different amitraz concentrations used (0.1%, 0.4% and 4%) caused fibrillae erection in at least 85% of mosquitoes. This effect was maintained for 30 min, and after 60 min still round 75% of mosquitoes keep expanded fibrillae. Twenty-four hours after exposure, the effect had greatly waned, but approximately 20% of mosquitoes still exhibited extended antennal fibrillae. **b)** Knockdown assay after amitraz exposure. Amitraz does not cause mosquito knockdown during an observation period of one hour post exposure. (**a, b**) Each category represents the mean of 3 replicates using 5 mosquitoes. **c)** Effects of amitraz on mosquito survival. Kaplan-Meier survival curve shows no differences in survival between control and 0.4% amitraz treated mosquitoes (Mantel–Cox test, p>0.05). Each category represents 3 replicates using 25 mosquitoes each. **d)** Diagram showing (created using BioRender.com.) the methodology used to assess the effects of amitraz exposure on mosquito mating behaviour. **e)** Mating behaviour assays in small laboratory cages. Treatment cages, where both male and female mosquitoes were exposed to 0.4% amitraz, show a reduction in insemination rates compared to controls (GLM (binomial), p<0.01). **f**) Mating behaviour assays in large laboratory cages replicate results of small cages, as treatment cages (both males and females exposed to 0.4% amitraz) showed a reduction of female insemination rates (GLM (binomial) p<0.01). **g**) Mating behaviour assays performed in semi-field facilities in Ifakara (Tanzania). In this case, only male mosquitoes were exposed to 4% amitraz. No differences in mating rates were observed for amitraz-treated males in natural conditions (GLM (binomial) p>0.05). Each dot represents the mean of a replicate using 50 (a), 250 (b) and 50 (c) females in the initial crosses, total of 4 (a), 4 (b) and 7 (c) replicates. **h**) Effects of amitraz exposure on numbers of eggs laid and larvae. Females were exposed to 0.4% amitraz twice before blood-feeding and before egg laying. No changes were observed in the eggs numbers (t-test, p>0.05) or hatching rates (GLM (binomial) p>0.05) between control and treatment cages, indicating that amitraz exposure does not affect female fertility.

We then tested the effects of 0.4% amitraz exposure on mosquito mating behaviour. Males and females were exposed and allowed to mate for two swarming periods in small laboratory cages (Fig. 5e). Insemination rates were reduced in males exposed to amitraz compared with controls (control: 44.5 ± 3.7%, amitraz: 28.9 ± 3.3%, GLM (binomial) p<0.01). We replicated the experiments in large laboratory cages (Fig. 5f), where mosquitoes were observed to form proper swarms (*40*) and noted a similar reduction in insemination rates (control: 53.0 ± 3.4%, amitraz: 37.8 ± 3.4%, GLM (binomial) p<0.01).

We then assessed the effect of amitraz on mating behaviour in a semi-field enclosure in a malaria endemic setting (Ifakara, Tanzania). We used the commercially available insecticide amitraz (Tiktit® 12.5% EC) and dissolved it to 4% following manufacturer’s instructions. Two-hundred males were exposed to 4% amitraz and placed in the semi-field chamber together with 50 non-exposed females (Fig. 5g). After one swarming period, the insemination status of females was assessed. We did not observe significant differences in insemination rates of females crossed to control and treated males (control: 24.9% ± 15.7%, amitraz: 19.8 ± 9.5%, GLM (binomial) p>0.05), suggesting that despite the effects observed in the laboratory, amitraz does not significantly disrupt mosquito mating behaviour in semi-field conditions.

Lastly, we also wanted to explore whether amitraz could impair female fertility. We exposed mated females to 0.4% amitraz before blood-feeding and egg laying, then counted the number of eggs and their hatching rate (Fig. 5h). We did not find any difference between control and treatment females (hatching rate: control: 85 ± 0.4%, amitraz: 83 ± 14%), indicating that amitraz exposure does not affect female fertility.

### AgOctβ2 sensitivity to amitraz is equivalent to its honeybee ortholog

Octβ2R has been shown to present different sensitivity to amitraz across arthropods (*30*). To assess the sensitivity of AgOctβ2, we studied its responses in cell culture (*41*). We compared these to two orthologues that have well characterised amitraz responses (*30*): the susceptible mite *Varroa destructor* (VdOctβ2) and the resistant honeybee *Apis mellifera* (AmOctβ2). We developed a cell-based system to express Octβ2R orthologues in HeLa cells co-expressing the cAMP biosensor G-Flamp2 (*42*) and monitored receptor activity in response to varying concentrations of ligand (Fig. 6a).

**Figure 6:**
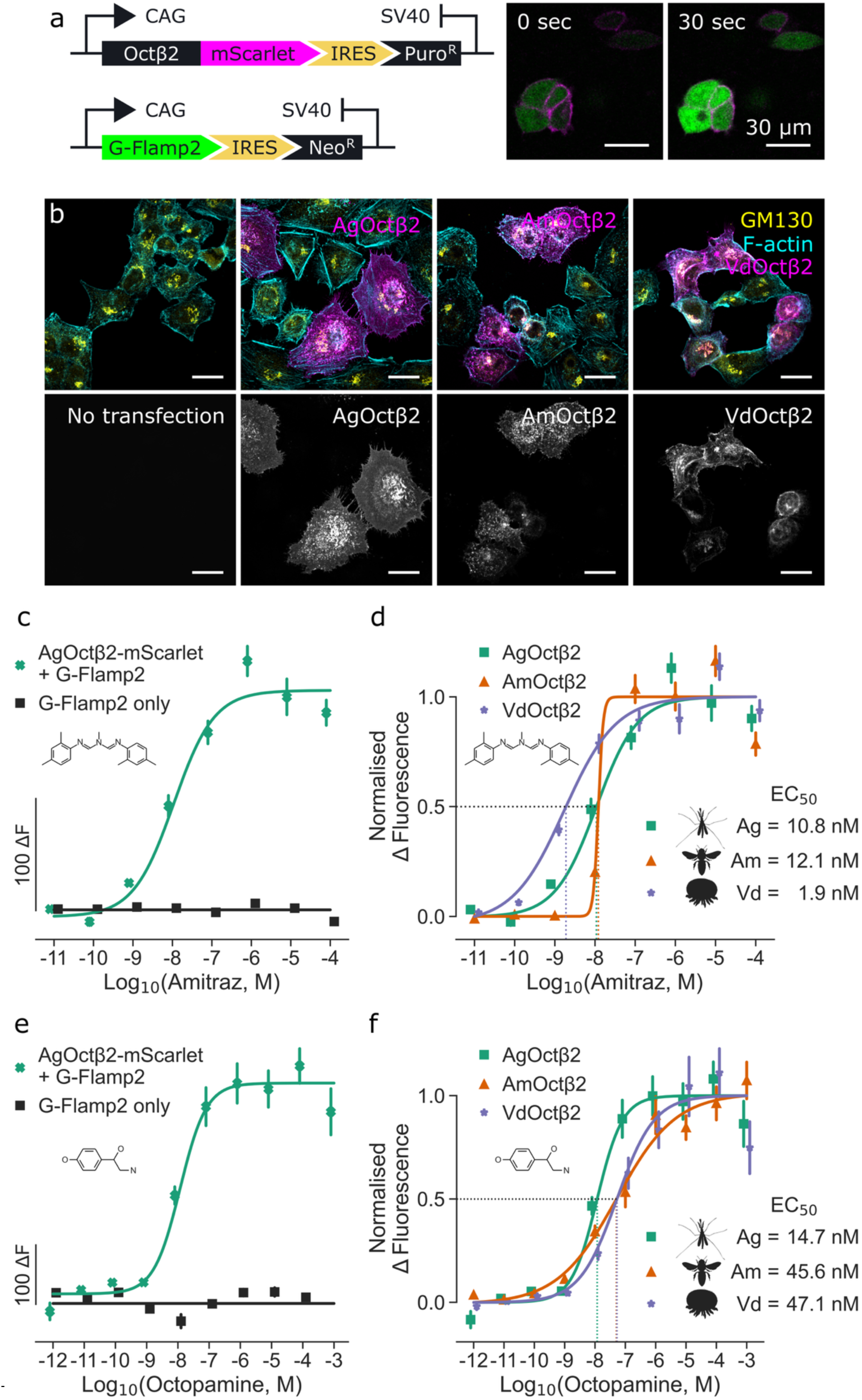
AgOctβ2R shows lower sensitivity to amitraz than the mite ortholog, but similar to the honeybee receptor. **a)** Confocal images of cell lines. Top row shows mScarlet-tagged receptor (magenta), Phalloidin staining F-actin (cyan) and αGM130 at the Golgi (yellow). Bottom row shows mScarlet channel only. Scale bars represent 10 µm. **b)** Schematic of constructs used. In the top construct, a CAG promoter drives expression of the Octβ2 orthologue, which is tagged at the C-terminus with mScarlet. In the bottom construct, the same promoter drives expression of cAMP-responsive G-FLAMP2 in the cytoplasm. Each construct was integrated using Tol2 transposase. Distinct antibiotic markers allowed both to be integrated in the same cells. **c)** Example frames showing increased G-FLAMP2 intensity after the addition of amitraz. **d)** The raw change in fluorescence before and after the addition of amitraz in G-FLAMP2 cells expressing Octβ2 from *An. gambiae* (green crosses) compared to those with no receptor (black squares). Points show mean and standard error. **e)** Normalised change in fluorescence comparing responses to amitraz in G-FLAMP2 cells expressing Octβ2 from *An. gambiae* (green squares), *Apis mellifera* (AmOctβ2, orange triangles) and *Varroa destructor* (VdOctβ2, purple stars). Dotted lines show where EC50 values were interpolated. **f)** The raw change in fluorescence before and after the addition of octopamine in G-FLAMP2 cells expressing AgOctβ2 (green crosses) compared to those with no receptor (black squares). Points show mean and standard error. **g)** Normalised change in fluorescence comparing responses to octopamine in G-FLAMP2 cells expressing AgOctβ2 (green squares), AmOctβ2 (orange triangles) and VdOctβ2 (purple stars). Dotted lines show where EC50 values were interpolated.

We first confirmed membrane expression of each orthologue using a C-terminally mScarlet. To do this, we imaged transiently transfected cells co-stained with the Golgi marker GM130. All cell lines showed good membrane localisation (Fig. 6b). We then analysed amitraz responses in cell lines expressing AgOctβ2, AmOctβ2 and VdOctβ2. To confirm HeLa cells do not respond to amitraz via an endogenous mechanism, we compared the fluorescent response (fluorescence after drug minus before; ΔF) of cells expressing both AgOctβ2 and G-Flamp2 to cells expressing only G-Flamp2 (Fig. 6c,e). Whilst the cells expressing the heterologous receptor showed a clear dose-response, G-Flamp2-only cells showed no effect of increasing amitraz doses. We then compared the response of lines expressing each receptor, by normalising each dataset to the maxima and minima of their respective logistic fits. We found that the mosquito receptor was 5.7-fold less sensitive than the mite receptor (Fig. 6d, EC_50_AgOctβ2= 10.8 nM, EC_50_VdOctβ2= 1.9 nM), but had comparable sensitivity to the honeybee receptor (EC_50_AmOctβ2= 12.1 nM; 1.1-fold compared to AgOctβ2). Therefore, in our heterologous system, the mosquito receptor shows a similar sensitivity to amitraz as honeybee – a species known to be resistant to hyperactivation. We also analysed the receptors’ sensitivity to octopamine, the native ligand, and found that AgOctβ2 is approximately 3-fold more sensitive than the mite and honeybee receptors (Fig. 6f, EC_50_AgOctβ2= 14.7 nM, EC_50_AgOctβ2= 45.6 nM, EC_50_VdOctβ2= 47.1 nM).

### AgOctβ2 residues can be substituted to alter the receptor sensitivity to amitraz

To identify differences in the molecular structure of the three receptors that could explain their different sensitivity to amitraz, we compared their amino acid sequences in the region around the predicted ligand binding pocket (Fig. 7a). We used AlphaFold structures to conduct molecular docking studies of octopamine, amitraz and DPMF (amitraz’s main metabolite). Each ligand was docked ten times using AutoDock Vina, and the number of times each residue was within 5 Ångstroms of the ligand was recorded. This allowed us to build a profile of residues with potential involvement in ligand binding across receptor orthologues and ligands (Fig. 7b).

**Figure 7:**
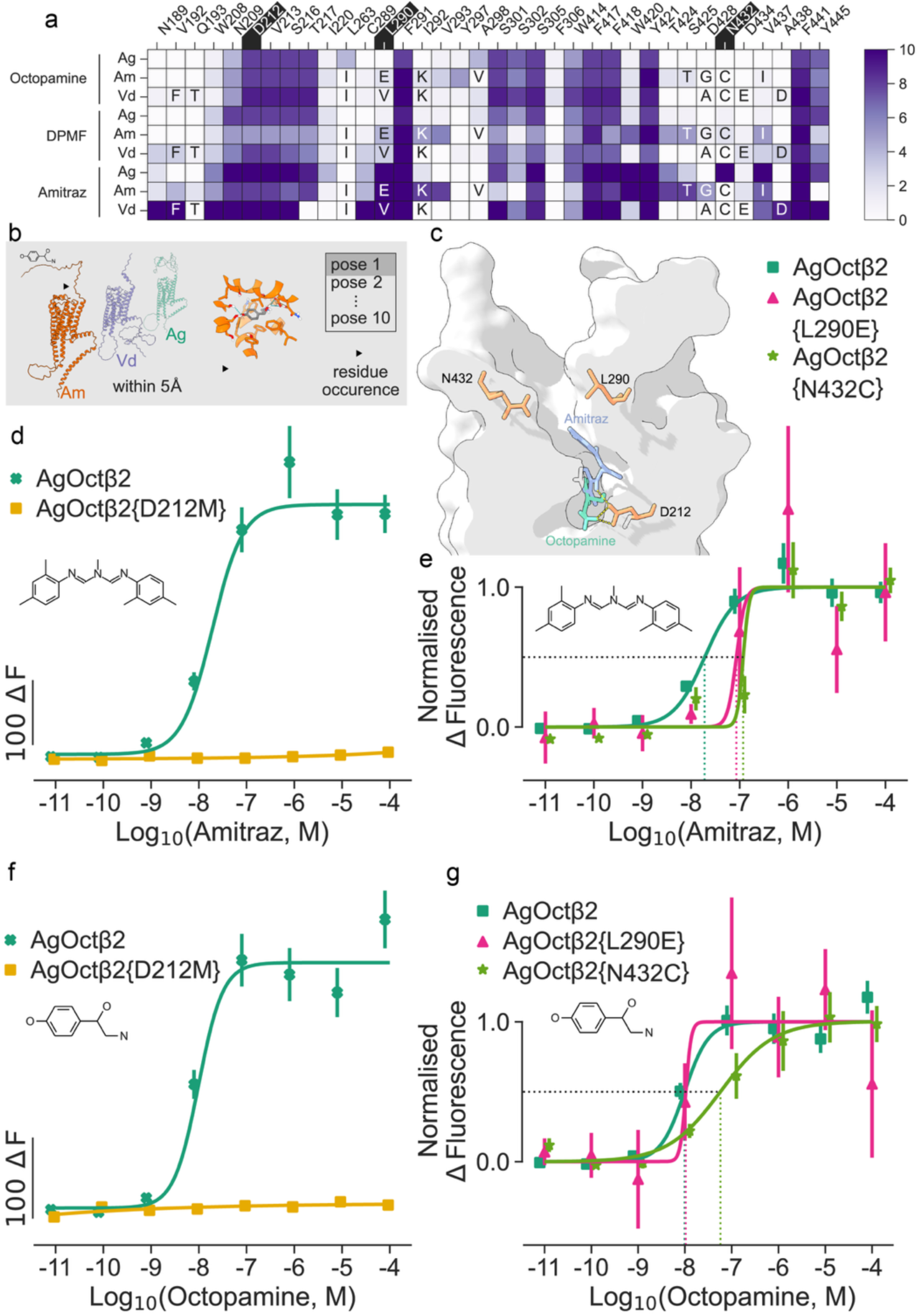
Amino acid substitution studies reveal AgOctβ2 residues involved in amitraz binding. **a)** Heatmap showing the occurrence of residues in repeats of each docking experiment for mosquito, honeybee and mite receptor. Each molecule was docked into each receptor ten times. There is one row for each molecule-receptor docking experiment, and one column for each residue (occurring >3 times). The colour of a square represents the number of times each residue was within 5 Angstrom of a pose. **b)** Summary of the above experiment. **c)** Cross-section of mosquito receptor with octopamine and amitraz docked in the binding pocket. Residues studied in red; octopamine in green; amitraz in purple; Hydrogen bonds at D212 in yellow. **d)** Raw change in fluorescence from before to after addition of drug (ΔF) for wt receptor (AgOctβ2) and D212M mutant at various doses of amitraz. **e)** Normalised change in fluorescence (ΔF normalised to minimum and maximum of four parameter logistic fit) for wt receptor, L290E and N432C mutants, at various concentrations of amitraz. **f)** Raw change in fluorescence from before to after addition of drug (ΔF) for wt receptor and D212M mutant at various doses of octopamine. **g)** Normalised change in fluorescence (ΔF normalised to minimum and maximum of four parameter logistic fit) for wt receptor, L290E and N432C mutants, at various concentrations of octopamine.

Some residues were predicted to be part of the binding pocket of all three ligands and were conserved in all three species, while others appeared to be ligand-specific or species-specific. We selected three residues that we considered of particular interest for their potential role in amitraz binding in mosquitoes (Fig. 7c, D212, L290 and N432). In other systems, residue D212 (located in the third transmembrane domain) has been implicated in ligand binding (*37*, *43*). This residue was conserved across the three species and predicted to be in the binding pocket of all three ligands: octopamine, amitraz and DPMF (Fig. 7a,c). To confirm its involvement in ligand-binding, we replaced the polar D212 of AgOctβ2, with a methionine which is non-polar and hydrophobic. We observed that mutating this residue completely ablates binding of both octopamine and amitraz (Fig. 7d, f), confirming that D212 is integral to ligand binding in *Anopheles*.

Residue L290 was found in the neighbourhood of docked amitraz, but not octopamine or DPMF (Fig. 7a). A previous study has shown that this residue, plays an important role in the increased amitraz sensitivity of the mite receptor relative to the honeybee(*30*). Although this residue is different in all three species, the mosquito (leucine) and mite (valine) are both hydrophobic branched-chain amino acids. By contrast, the honeybee residue (glutamic acid) is hydrophilic.

We therefore speculated that this residue could promote sensitivity in mosquitoes. We mutated the mosquito residue to the corresponding amino acid in honeybees and examined the pharmacological impact. In support of our hypothesis, AgOctβ2^L290E^ showed reduced sensitivity to amitraz (Fig. 7e, AgOctβ2 EC_50_= 19.1 nM; AgOctβ2^L290E^ EC_50_= 85.4 nM). However, the receptor showed un-altered sensitivity to octopamine (Fig. 7g, AgOctβ2 EC_50_= 9.84 nM; AgOctβ2^L290E^ EC_50_= 10.4 mM), supporting the docking experiments that implicate this residue in amitraz binding only. Finally, we selected residue N432 as it was conserved in mites and honeybees (cysteine) but not in mosquitoes (asparagine). It was not predicted to be involved in binding octopamine or DPMF in any species, and only proximal to amitraz when docked in mosquitoes. We replaced the mosquito asparagine with the corresponding residue in honeybees and mites (cysteine) and found that the resulting AgOctβ2^N432C^ showed reduced responses to both amitraz (Fig. 7e, EC_50_ = 118 nM) and octopamine (Fig. 7g, EC_50_ = 57.3 nM). This residue is far from the predicted binding site of the different ligands (Fig. 7c) suggesting this mutation may have reduced access to the pocket for all ligands.

## Discussion

Mosquito mating and reproduction are clear targets to reduce *An. gambiae* numbers. In this paper, we explored this approach at multiple levels. Firstly, we demonstrate that octopamine modulates different aspects of male audition via AgOctβ2. Moreover, *AgOctβ2* mutant males exhibited a severe reduction of insemination rates, likely because of its auditory defects. Additionally, we showed that mutant females are sterile through an unrelated biological mechanism. These phenotypes suggest that AgOctβ2 is an ideal target for malaria reproductive control. To explore this, we developed novel screening approaches and tested the ability of various compounds, including the insecticide and *AgOctβ2* agonist, amitraz, to disrupt mating. We showed that amitraz exposure causes subtle reductions in insemination rates, but no effects in female fertility, ruling out its application to disrupt reproduction more widely. We then conducted molecular docking and cell-based assays across different arthropods to identify putative AgOctβ2 residues involved in amitraz binding in mosquitoes, aiming at aiding future compound design. Our work points to AgOctβ2 as a promising target to disrupt mosquito reproduction, although it also highlights the challenges related to applying this approach for malaria control in the field.

### Disrupting AgOctβ2 signalling in *An. gambiae* severely impairs mosquito audition and mating

Audition in males drives the detection of the female mating partner during courtship (*44*, *45*). We have previously proposed that hearing offers valuable targets for mosquito control (*18–20*, *22*, *46*). The mosquito auditory system is highly complex (*47–50*) and offers multiple opportunities to disrupt its function. A recent study demonstrated that knock out of the TRPV channel *trpVa,* the homolog of the *Drosophila* channel *Iav*, in *Ae. aegypti* males caused deafness and inability to mate with females (*21*). In malaria mosquitoes, we previously showed that octopamine modulates mechanical aspects of auditory function through AgOctβ2 activation (*22*). Mutant males do not respond to octopamine, and consequently are unable to erect their fibrillae at swarm time and show defects in their auditory tuning.

To further explore the auditory role of octopamine, we used frequency-modulated sweeps to extract auditory tuning frequencies (best frequencies), as they produce responses of maximal amplitude. The male mosquito ear presents different mechanical and electrical best frequencies as a result of their distortion product-based hearing (*17*). We confirmed previous results (*22*) that octopamine injections caused a shift in the mechanical tuning to higher frequencies in males (∼ 180 Hz higher, corresponding to a 1.49 increase *f_2_/f_1_*), but not females (Fig. 1e). Interestingly, a previous publication studying octopamine’s role in *Cx. quinquefasciatus* mosquitoes showed the same increase of 1.49 times following octopamine injection (*50*). This striking similarity points at a conservation of octopamine’s auditory role across mosquito species, although the degree of sexual dimorphism varies. Recent studies in *Culex pipiens* and *Ae. aegypti* reported effects of octopamine in females (*51*, *52*) while we find no evidence of this in *An. gambiae*. These species-specific functional variations likely arise from neuroanatomical differences in the efferent innervation across females from different species (*19*, *46*).

We also found that octopamine increases the electrical tuning frequency in males (Fig. 1f, ∼ 90 Hz higher, corresponding to a 1.42 increase *f_2_/f_1_*). The ratio between pre-injection and post-injection best frequencies is thus similar for the mechanical (1.49 ratio) and electrical (1.42) tuning frequencies, suggesting that their mechanism of action may have the same origin. We also observed that octopamine enhances the electrical responses of auditory neurons to mechanical stimuli. By applying force-steps and simultaneously recording flagellar displacements and CAP responses, we show that CAP amplitude strongly increased relative to flagellar displacements upon octopamine injections (Fig. 1h). Similar effects of octopamine have been previously described in mechanosensory organs of the spider *Cupiennius salei* (*53*). These results illustrate that octopamine signalling through AgOctβ2 is a master modulator of auditory function in the swarm(*22*), as it induces the erection of the antennal fibrillae and controls flagellar stiffness, mechanical and electrical tuning, and auditory sensitivity. Although many of these effects might be interconnected, the complex pattern of octopamine innervation in the JO of other mosquito species at different subcellular localizations of the auditory neuron (*19*, *50*) suggest that it could modulate these mechanisms independently, raising interesting questions about how octopamine enables auditory plasticity.

We further examined potential phenotypes of *AgOctβ2* knockouts in auditory behaviours. We analysed phonotactic responses of males and observed that knockout mosquitoes were less attracted to sound than wild-types (Fig. 2a). As attraction to the flying female is an essential component of mosquito courtship (*12*), we studied the ability of mutant mosquitoes to mate. We crossed wild-type and knockout male and female mosquitoes in different combinations and observed that mutant males were less able to mate (up to ∼ 90%) independent of the female genotype. To our knowledge, this is the first time that the genetic disruption of hearing in malaria mosquitoes is shown to affect mating behaviour. Importantly, mutant mosquitoes are not strictly deaf, as they can respond to auditory stimulation, but they exhibit altered responses to octopamine, affecting its auditory function (*22*). We hypothesize that the lack of fibrillae erection, altered auditory tuning and reduced auditory sensitivity caused by impaired octopamine signalling hinders the detection of mating partners. Our results also stress the highly dimorphic nature of mosquito mating, and the predominant role of the male in female acoustic detection, suggesting that potential vector control tools aimed at disrupting mosquito audition should be focussed on males.

### Knocking out *AgOctβ2* causes female sterility, suggesting novel applications for vector control

Beyond the auditory defects displayed by *AgOctβ2* knockout males, we found that the mutation causes female sterility. Octopamine has been extensively reported to be involved in reproduction across insects (*37*, *54–58*), and different octopamine receptors are expressed in the female reproductive tract, including *AgOctβ2* orthologs (*36*, *59*, *60*). Indeed, *Drosophila melanogaster Octβ2* knockout females are also sterile due to ovulation defects that cause an inability to lay eggs (*35*). Distinct from this phenotype in *Drosophila*, we did not observe a decrease in the number of eggs laid by mutant females (Fig. 2f). However, all eggs laid by *AgOctβ2* knockout females failed to hatch (Fig. 2h), implying female sterility. These results suggest that eggs might not be fertilized by sperm. Octopamine has been shown to regulate the release of sperm from storage in the female reproductive tract of *Drosophila* before egg fertilization (*61*, *62*). It is plausible that female sterility in our mutant is caused by defects in sperm release, but further investigation is needed to confirm this hypothesis.

From a vector control perspective, the female sterility phenotype and the mating defects in mutant males turn AgOctβ2 into a novel target to control mosquito reproduction. The development of reproductive control strategies for malaria mosquitoes has been mainly explored by generating genetic modifications that cause female sterility and are spread into mosquito populations using gene drives (*9–11*). Indeed, female sterility phenotypes observed in *AgOctβ2^-^* suggest it as a promising candidate gene for this technology. However, the reduced mating ability of mutant males might hinder its applicability as gene drive target, as this phenotype would limit the spread of mutant alleles in the population. Conversely, the phenotype offers opportunities to explore the development of pharmacological approaches to disrupt reproduction and their knock-on effects on mosquito population dynamics and disease transmission.

### Amitraz cannot be directly applied to disrupt mosquito reproductive behaviour

To examine if the pharmacological disruption of AgOctβ2-signalling would affect male mating ability or female sterility, we first screened different compounds that have been reported to antagonize Octβ2 receptors. The control of the antennal fibrillae erection pattern by AgOctβ2 (*22*) offers an elegant system to test drug effects *in vivo*. We applied different delivery methods (including injections, topical exposure and feeding) to administer chemical compounds. Although performing injections requires some technical equipment, feeding and topical exposure procedures are simple methods that can be used in low-resource settings. In our laboratory conditions, we also designed a high-throughput video-based feeding system to track mosquito fibrillae status in response to fed compounds. The system allows the testing of multiple compounds or doses at the same time (Fig. 3).

We chose mianserin and epinastine for further behavioural tests due to their higher *in vivo* potency as AgOctβ2 antagonists in the feeding assay. However, feeding both epinastine or mianserin caused very high mortality in males (Suppl. Fig. 3), making it difficult to interpret the effects on mating behaviour effects (Fig. 3d). Even so, neither drug seemed to inhibit male mating capacity.

We then assayed mating effects of the insecticide amitraz, which activates AgOctβ2 *in vivo* (*22*), and is widely used for mite and tick control. We previously showed that amitraz exerts auditory effects on male mosquitoes by inducing antennal fibrillae erection (*22*). Interestingly, honeybees exposed to amitraz present higher mating frequencies than control individuals (*63*). We investigated the behavioural implications of amitraz’s auditory effects and showed that exposing males to 0.4% amitraz causes a 34% and 29% reduction in female insemination rates in small and large laboratory cages, respectively (Fig. 5e,f). We further conducted mating behavioural studies in a semi-field system in a malaria endemic setting using commercially available insecticide. In the semi-field, we could not replicate amitraz’s effects on mating behaviour, and mating rates were not statistically reduced (Fig. 5c). This could be due to the different mosquito strains used – lab-adapted G3 compared to field-derived *Anopheles gambiae* s.s (lab adaptation has been found to reduce mosquito mating competitiveness (*64*)), or different effects of the pure compound vs insecticide formulations (*65*). In any case, the results in the semi-field rule out the application of amitraz as a mating disruptor. Moreover, amitraz exposure did not cause any effect on female sterility (Fig. 5h), excluding potential applications to disrupt mosquito reproduction and suggesting that new chemistries with higher potency or with antagonistic effects on AgOctβ2 are needed.

### Development of new tools to target AgOctβ2 pharmacologically

Previous studies have shown that the sensitivity of Octβ2 receptors to amitraz varies across arthropods (*30*). This selectivity has provided an opportunity to target parasitic mites, reducing off-target effects in insect pollinators. We first aimed to test the potency of amitraz on AgOctβ2. The fact that amitraz causes low mortality in *An. gambiae* (compared to other arthropods) suggests lower sensitivity of the receptor. Using cAMP-reporter assays, we demonstrated that AgOctβ2 sensitivity to amitraz is reduced compared to the mite, *Varroa destructor*, receptor, more similar in profile to that of the honeybee, *Apis mellifera*. The EC_50_ values of the three receptors are very similar for octopamine.

To identify residues of importance for future molecule design in the mosquito receptor, we compared its ligand docking profile to the amitraz-susceptible mite and the amitraz-resistant honeybee. Firstly, we show that L290 within the pocket is important for the, albeit low, susceptibility of AgOctβ2 to amitraz. This residue appears to show diversity across taxa and is orthologous to a valine in *Varroa* (that is also important for susceptibility(*30*)), and a glutamic acid in *Apis*. Such non-conserved residues that are still functionally important could provide good candidates for molecule design, as they have potential to increase specificity and reduce off-target effects in other organisms. Another such residue is N432 identified near the mouth of the binding pocket. Both other organisms have a cysteine at this position, and when we mutated the *Anopheles* asparagine to mimic this, the receptor became less susceptible to both amitraz and octopamine. This residue could provide a good target for antagonists aiming to block the mouth of the binding pocket.

### Future directions

Our study presents the role of *AgOctβ2* in *An. gambiae* mating and fertility. The phenotypes reported suggest AgOctβ2 as a novel and promising target to control malaria mosquito reproduction. The fact that AgOctβ2 is already targeted by the pesticide amitraz, which is widely used to control parasitic mites provides evidence of the applicability of this approach in agriculture and public health. Our study suggests that the structure of amitraz could be optimized to increase its potency against AgOctβ2. Given the demonstrated selectivity of amitraz to mite Octβ2 receptors, it might be plausible to modify its structure to selectively target mosquito receptors. This approach would allow for exploring new pathways for malaria mosquito control based not on killing mosquitoes, but rather on modifying their mating behaviour and reproduction to reduce vectorial capacity. These “mating disruptors” could be delivered using attractive sugar baits or even included in insecticide-treated net formulations (*66*). Further studies are required to identify lead compounds and conduct field and modelling studies to endorse this approach as an alternative strategy for malaria control.

## Materials and Methods

### Mosquito rearing and entrainment

*An. gambiae* G3 strain (wild-type) and *AgOctβ2* knockout mosquitoes were reared in our insectary at UCL Ear Institute. All *AgOctβ2^-^* mosquitoes used were homozygous. Experimental pupae were sex-separated and males and females emerged in different cages. Mosquitoes were then circadian entrained for at least three days using environmental incubators (Percival I-30 VL, CLF PlantClimatics GmbH) to 12 h: 12 h light/dark cycle (maximum light intensity was 80 μmolm-2 s-1 from 4 fluorescent lamps) with 1-hour ramped light transition to simulate dawn and dusk (ZT0-ZT1 and ZT12-ZT13, respectively; ZTX is the formalized notation of an entrained circadian cycle’s phase), 28 °C and 80% humidity. All experiments were conducted with 3-to-7-day old mosquitoes.

### Compound delivery

Compounds used were octopamine hydrochloride (Generon, CAS 770-05-8), phentolamine hydrochloride (Sigma-Aldrich, P7547), epinastine hydrochloride (Sigma-Aldrich, E5156), mianserin hydrochloride (Sigma-Aldrich, M2525) and amitraz (Sigma-Aldrich, 45323). Solvents and delivery methods varied (Table 2). Solutions were prepared fresh each day.

**Table 1:**
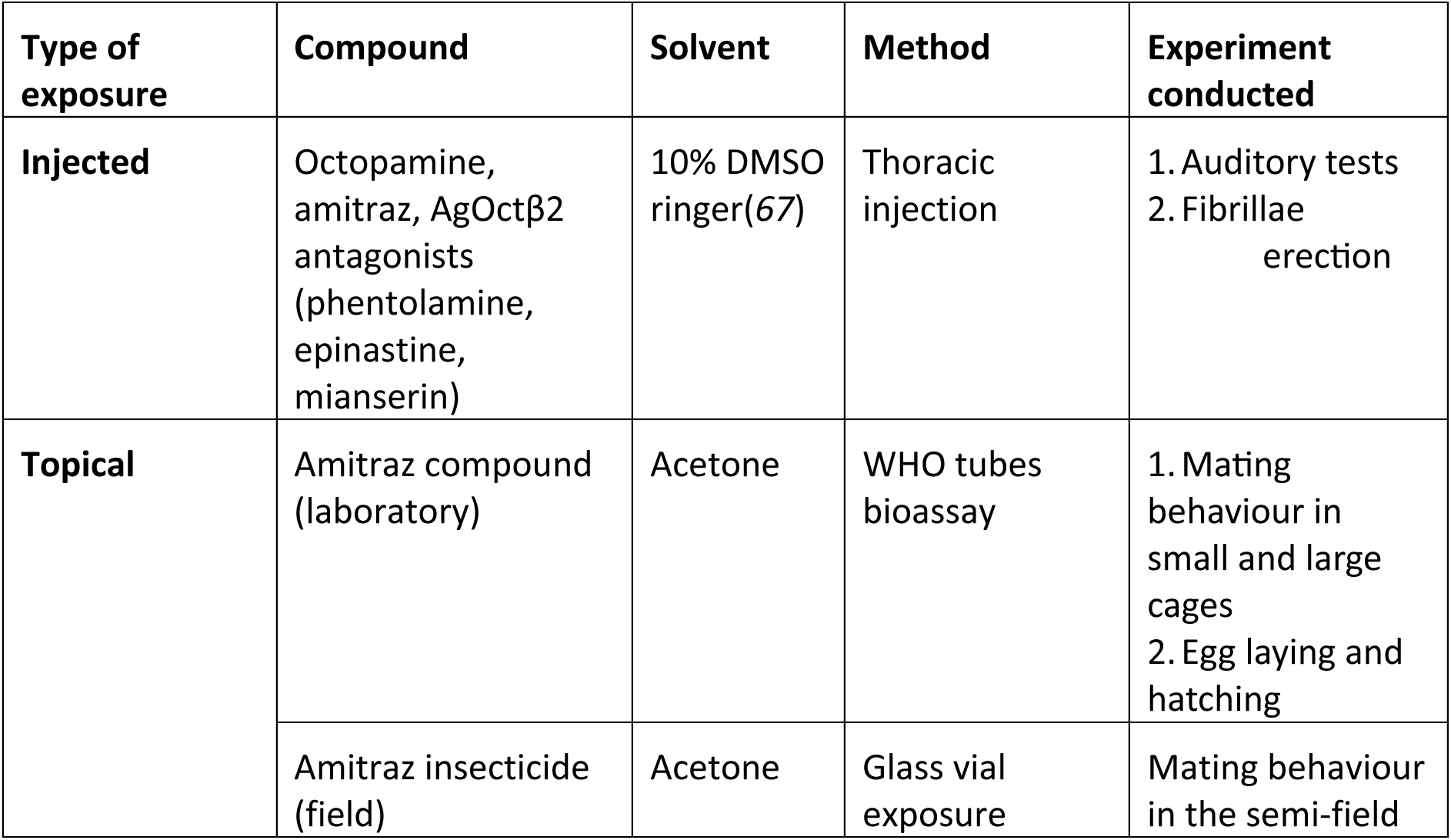
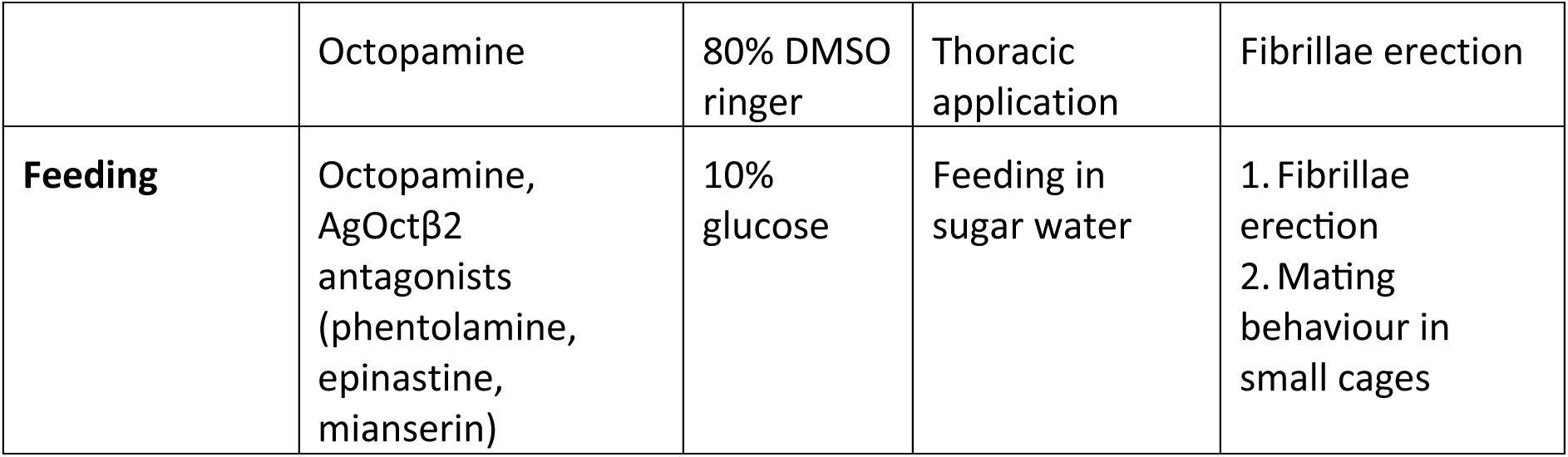
Summary table of different compound delivery methods and experiments conducted. Table 1: Summary of methods used to deliver compounds, including the compounds, delivery methods, and the experiments that were performed. Details of the experimental procedures can be found in the Methods section.

### Compound injection

Mosquitoes were anesthetized on ice and immobilised to a plastic rod using blue-light-cured dental glue, except for both antenna (in fibrillae erection experiments) or the right flagellum (for auditory tests, in this case, also the pedicel to head joint was immobilised, see below) (*46*). Borosilicate microcapillaries (Drummond) were pulled with a p-1000 Sutter micropipette puller. A Nanoject III was used to inject 200 μl of compound into the thorax (Table 2). Compound effects were tested immediately after either assessing the fibrillae erection pattern or conducting auditory tests (see below).

### Compound topical exposure

#### WHO tube bioassays

WHO insecticide exposure tubes were used to expose mosquitoes following WHO tube bioassay procedures (<u>link</u>). Acetone (control) or 0.4% amitraz dissolved in acetone were applied to 12 x 15 cm pieces of aluminium foil and dried in a fume hood. The aluminium foil was placed inside the WHO tube to cover its surface. Twenty-five mosquitoes were introduced to the tube and exposed for 3 minutes. Upon exposure, mosquitoes were transferred back to paper cups and provided with 10% glucose solution for further experiments.

#### Glass vial exposure

A stock solution of insecticide amitraz (Tiktit® 12.5% EC) was dissolved in acetone to 4 % following manufacturer instructions. A 20 ml glass bottle was coated with either control (acetone) or treatment solutions and allowed to dry for 2 hours in a dark and cool room. Five 3-days old *An. gambiae s.s* mosquito males were aspirated into the glass vials and exposed for 3 min. Upon exposure, mosquitoes were transferred back to the paper cups and provided with 10% glucose solution for further experiments.

#### Thoracic application

Mosquitoes were glued to a plastic rod as explained in “Compound injection”. Compounds were dissolved in 80% DMSO to facilitate cuticle penetration. A pipette was used to apply a 0.5 µl drop of the solution to the mosquito thorax. Effects of the compounds on fibrillae erection were assessed immediately after.

### Compound feeding

#### Video recorded feeding assay

Plates were 3D-printed with PLA using an Ultimaker 3 (STL file provided as Suppl. Fig. 4). Epoxy resin (Araldite) was used to adhere the bottom of the plate to a piece of clear cast acrylic. Nylon M3 bolts were also attached to the plate with epoxy resin, facing up through the printed holes. These aligned with holes made using a push drill, in two sections of clear-cast acrylic that were cut to cover the wells. These pieces of acrylic lay flush with sugar-soaked dental swabs (1 cm diameter; ± compounds) arrayed in the half-pipe sections to the side of each well. M3 nuts were then used to secure the top pieces of acrylic during experiments. To capture images, a Raspberry Pi HQ camera with the infrared filter removed, paired with a wide-angle lens, was attached to the ceiling of an incubator. A Raspberry Pi 3 was then used to capture a frame every 30 seconds. The setup was illuminated from below using strips of infrared LEDs which were diffused using opal acrylic.

Three-to-5-day old male mosquitoes were sugar starved in separate cages for 16 hours followed by water starvation for 3 hours. Mosquitoes were then anesthetized on ice, transferred to a plate and imaged over 90 minutes. Antagonists (5 mM mianserin, 10 mM epinastine and 20 mM phentolamine) were dissolved in 10% glucose, along with 36.7 mM octopamine.

Google Teachable Machine was used to train an image classifier (MobileNet) using 756 images of mosquitoes with erect antennal fibrillae, 763 collapsed images and 662 empty images. Half of each image set was generated using augmentation (Albumentations package in Python (*68*)), with random cropping, square symmetry, sharpening and blurring. The following parameters were used during training: 1000 epochs, batch size of 32, learning rate of 1 x 10^-6^. The training and validation loss curves showed negligible signs of overfitting or underfitting. A Python-based image analysis pipeline was established using OpenCV, whereby videos were opened frame-by-frame, wells were identified by performing contour detection (this was done on a thresholded and closed image, with contours filtered by size) wells were assigned IDs based on relative location, and the image classifier was used to predict the fibrillar state of a sub-image of each well. The final image classifier model and image segmentation pipeline are available on GitHub (https://github.com/mosquitome/buzz-feeder).

#### Fibrillae erection assessment

Experiments were performed at ZT4 to avoid any endogenous fibrillae erection that occurs at swarm time. Compounds were either injected, topically applied or fed (see above). Different antagonist concentrations (0.1 mM, 1 mM, and 10 mM) were administered together with 1 mM octopamine. Five mosquitoes were tested for each concentration and the proportion erecting their fibrillae was visually assessed over a 30-minute window immediately after exposure. Each experiment was repeated five times.

For amitraz exposure experiments using WHO test tubes, fibrillae erection was observed for 72 hours, and the fibrillae status was observed 5, 10, 30, 60 min, 24, 48 and 72 hours after exposure.

#### Knockdown assays

Mosquitoes were exposed to 0.1%, 0.4% and 1% amitraz following WHO tube bioassay procedures. Mosquitoes were visually monitored for knockdown effects (paralysis) for 1 hour; at 5, 10, 30 and 60 min.

#### Lethality experiments

Twenty-five two-to-three-day old female mosquitoes were exposed in WHO tubes bioassays to 0.4% amitraz. Females were then released into small cages (15 x 15 cm) and mortality was recorded daily.

### Tests of auditory function

#### Laser-Doppler vibrometry (LDV) and electrophysiology

Glass vials containing five mosquitoes were removed from incubators at ZT12 and anaesthetised on ice. They were then mounted on blue tac at the end of plastic rods using curable dental glue as previously described (*22*, *46*). The mosquito body was immobilised with glue to minimise disturbances caused by mosquito movements but leaving thoracic spiracles free for the mosquito to breathe. The left flagellum was glued to the head and further glue was applied between the pedicels, with only the right flagellum remaining free to move. All measurements were taken in a temperature-controlled room (22°C) at ZT12-ZT14.

Following this gluing procedure, the rod holding the mosquito was placed in a micromanipulator on a vibration isolation table, with the mosquito facing the Laser-Doppler vibrometer at a 90° angle. The laser was focused at the second flagellomere from the flagellum tip. Two electrostatic actuators were positioned symmetrically either side of the flagellar tip to allow for push- and pull-electrostatic flagellar actuation. For electrophysiological recordings, a reference electrode was inserted in the mosquito thorax, and a recording electrode in the antennal nerve. Force-step stimulation was used to calibrate the maximum flagellar displacement to approximately ± 8000 nm. All recordings were made using a PSV-400 Laser-Doppler Vibrometer (LDV, Polytec) with an OFV-70 close up unit and a DD-500 displacement decoder. Spike 2 version 10 was used for data collection.

#### Frequency-modulated sweep stimulation and analysis

Mosquitoes were presented with 1-second-long periods of electrostatic stimulation, interleaved by 1 second-long (Fig.1c). Stimulation consisted of an electrostatic waveform as driving force with constant amplitude that swept linearly from 0 Hz to 1000 Hz (forward sweep), or 1000 Hz to 0 Hz (backward sweep), each repeated 60 times.

Experimental data was first prepared in Spike2; sections of the recordings for which the laser quality was deemed insufficient to maintain proper focus were manually removed. This was done through a diagnostic channel which measures and records the laser backscatter of the LDV. These files were then exported for analysis in Matlab (Mathworks) using a custom script. A smooth function (time constant 0.00025s) was applied to the flagellar displacement data, and a DC remove (time constant 0.015s) to the nerve response data. To identify the frequency of stimulation which resulted in the largest mechanical response (i.e. the largest displacement of the flagellum) or the largest electrical response (i.e. the largest nerve response), we utilised the ‘findpeaks’ function to identify sequential peaks, then calculated the absolute difference in magnitude between these peaks. We then identified the time at which the largest change in response magnitude was identified, then defined the stimulus frequency at this time as the best mechanical/ electrical frequency, averaging forward and backward sweep stimuli.

#### Force-step stimulation and analysis

LDV measurements of flagellar displacements in response to electrostatic force-step stimuli of different magnitudes, as well as corresponding electrophysiological activity, were recorded simultaneously in Spike 2 (Fig. 1d). The maximum flagellar displacement elicited was 8000 nm, which was also used to calibrate experiments. Voltages of applied steps were also simultaneously recorded. Mosquito apparent flagellar mass estimates for data analysis were taken from (*46*). Force-step stimulation analysis was performed as in (*69*). Steady-state displacements were determined as the displacement of the flagellum prior to step off-set relative to the position prior to step onset. To compare nerve responses across individuals, CAP responses were normalised as ((V-V_max_)/(V_max_-V_min_)).

#### Phonotaxis assays

Twenty to 25 mosquitoes were entrained in a small cage (15 x 15 cm) for a minimum of 3 days before testing as explained above. At ZT13 (lights fully off, red lights on), the incubator door was opened, and randomised pure tones (250 – 700 Hz, 50Hz intervals) were played at one side of the cage and number of mosquitoes approaching the speaker were counted by the experimenter.

#### Mating experiments (laboratory small cages)

Before undergoing any of the experimental procedures explained below, male and female mosquitoes were entrained separately as detailed above. All mosquitoes were 3-to-7-day old when mating assays started.

#### Transgenic mosquitoes

For mating experiments using transgenic line *AgOctβ2*^-^, 50 males and 50 females in different combinations (*AgOctβ2*^-^ males and females, *AgOctβ2*^-^ males with wt females, wt males with *AgOctβ2*^-^ females and wt males and female) were introduced in a small laboratory cage (17.5 x 17.5 x 17.5 cm) and allowed to mate for three swarming periods (Fig. 2b). Females were collected in a paper cup and their insemination status evaluated by dissecting their spermathecae to ascertain the presence of sperm.

#### AgOctβ2 antagonist fed mosquitoes

50 male and 50 female mosquitoes were sugar starved in separate cages for 16 hours followed by water starvation for 3.5 hours (Fig.4a). The epinastine treatment was prepared by saturating cotton wool in a 35 mm x 10 mm petri dish with 5 mL of 2.5 mM epinastine or 5 mM mianserin diluted in 10% glucose solution. For the control, only 10% glucose solution was used. The treatments were added to the male and female cages separately at ZT10 on day 4. Prior to swarm time (between ZT11 and ZT12), males were added to female cages, with the number of dead or knockdown mosquitoes recorded. The treatments were replaced at ZT15, this time containing 7 mL drug in 10% glucose or 10% glucose only. Petri-dishes were covered in parafilm to prevent drying out of the cotton wool and a needle used to pierce holes in the film to ensure mosquito contact with the glucose solution. Control and treatment experiments were run in parallel. At ZT8 on day 5, numbers of dead or knockdown mosquitoes were noted. Females were collected, anesthetized on ice and the spermathecae were dissected.

#### Amitraz exposed mosquitoes

Male and female mosquitoes were exposed separately to 0.4% amitraz or control acetone in WHO insecticide exposure tubes 2-2.5 hours before swarm time (see “WHO tubes bioassays” section). Half an hour before swarm time, males and females were brought together in a single cage. Mosquitoes were allowed to mate for two swarming periods (Fig. 5d). Control and treatment experiments were run in parallel. Females were collected, anesthetized on ice and the spermathecae were dissected.

### Mating experiments (laboratory large cages)

#### Transgenic mosquitoes

Large experimental cages (5 x 1.25 x 2.60 m; L x W x H) constructed with wooden frames and enclosed in fine mesh were used for all experiments (Figure 2d). Each cage contained a 40 × 40 cm black-and-white square marker to stimulate swarming behavior, as previously described (*70*). Additional environmental features included a terracotta resting shelter maintained humid with a damp sponge and a black dome serving as a dry resting site. To promote swarming behavior, a simulated sunset was programmed to last 40 minutes. Ceiling lights gradually dimmed from full light to low light, followed by 30 minutes of twilight provided by horizon lights only. Mosquito pupae were separated by sex and allowed to emerge in individual BugDorm cages. On day 4 post-eclosion, 250 *AgOctβ2⁻* males and 250 females were released into each large cage three hours prior to the onset of simulated sunset. Two independent replicates were performed. Swarming behaviors was visually monitored on the day of release. The following day, female mosquitoes were collected using a Modified CDC Backpack Aspirator (John W. Hock Company, Gainesville, FL, USA), anesthetized on ice, and the spermathecae were dissected.

#### Amitraz-exposed mosquitoes

Two large cages (100 × 100 × 200 cm, L × W × H) constructed with wooden frames and enclosed in fine mesh were used for all experiments (Fig.5f). Each cage contained a black-and-white square marker (40 × 40 cm) placed at the center of the floor and black boards (40 cm high) lining three of the four walls. A light bulb positioned behind the rear wall simulated a horizon light source during the dark phase. A simulated sunset was programmed to last 40 minutes, consisting of a 30-minute gradual dimming of the ceiling lights followed by 10 minutes of steady dim red light, after which the lights were turned off completely. Horizon lights were manually turned on at the beginning of the sunset phase and remained on for 30 minutes after the transition to darkness.

Mosquitoes were provided with 10% glucose solution ad libitum. Pupae were sexed and allowed to emerge in separate cages. On day 4 post-eclosion, adult males and females were exposed to either 0.4% amitraz or an acetone control following the WHO tube bioassay protocol, in batches of 25 individuals. A total of 200 mosquitoes (100 males and 100 females) were released into the experimental cages 80 minutes before the simulated sunset. Swarming behavior was visually monitored on the day of release. The following day, mosquitoes were collected using a modified CDC Backpack Aspirator (John W. Hock Company, Gainesville, FL, USA). A subset of 100 females from each group was anesthetized on ice, and their spermathecae were dissected. The experiment was repeated four times.

#### Mating experiments (semi-field system)

These experiments were conducted in the semi-field system (SFS) chambers (*71*) at the Ifakara Health Institute (Tanzania), from 12 November to 03 December 2020, with parallel trials in paired control and treatment chambers (Fig.5a). Each SFS chamber was constructed with thick opaque white polyethylene to minimize external environmental variability, featuring a mesh density of 346 holes per square inch to allow light while preventing mosquito escape. A black cloth swarm marker (*72*), 1.5m x 1.5m was placed centrally on the floor of each chamber.

*Anopheles gambiae* s.s. mosquitoes were reared under standard insectary conditions (27 ± 2°C, 75% ± 10% relative humidity, 12:12 light:dark cycle) following MR4 guidelines (*73*). Larvae were fed Tetramin® fish food, and adults were maintained on 10% sucrose solution. Pupae were sexed, and males and females were kept separated to ensure virginity until experiments (*74*).

Male mosquitoes were exposed to either 4% amitraz or vehicle control (Glass vial exposure section) from 17:00–18:00 for acclimatization before release. At 18:00, 200 treated males and 50 unexposed virgin females were introduced into the chamber. Swarms were allowed to form, and collections were made between 18:30–18:50 (20 minutes post-swarm initiation) using a sweep net (*75*). Captured mosquitoes were aspirated into paper cups, and the following morning, females were frozen (−20°C for 5 minutes) and spermatheca dissected.

### Egg laying and hatching experiments (small cages)

#### Transgenic mosquitoes

Mosquitoes were allowed to mate for three swarming periods, see “Mating experiments (laboratory small cages)” section (Fig. 2e). Crosses included combinations of 50 males and 50 females of *AgOctβ2^-^* or wt mosquitoes. Female mosquitoes were collected, bloodfed and placed in individual cups. The number of eggs laid, and number hatched (1^st^ instar larvae) were calculated for each female.

#### Amitraz exposed mosquitoes

Fifty male and 30 female mosquitoes were allowed to mate for three swarming periods (Fig.4h). Females were then collected in a cage. The following day, females were exposed to 0.4% amitraz or vehicle following the WHO tubes bioassay protocol (see above). Four hours after exposure, females were blood fed and allowed to rest in cages for 3 days, when they were again exposed to 0.4% amitraz before egg laying. After this last exposure, control and treatment females were placed in cups. The number of eggs laid and hatched (1^st^ instar larvae) was counted.

#### Sequence comparison

The sequences of Octβ2 orthologues from *An. gambiae* (UniProt F5HKI1), *Ap. mellifera* (UniProt A0A7M7GMY8) and *V. destructor* (Uniprot A0A7M7KEY3) were aligned in Clustal Omega. Amino acids were numbered, and a reference table was built to compare any given residue in one species with the corresponding residues (or gaps) in the other two.

#### Molecular docking

Docking was performed using Alphafold structures of the orthologues described above (accessed 27/01/23) (*76*, *77*). SDF files of amitraz, octopamine and DPMF were downloaded from ZINC(*78*) and converted to PDB format in OpenBabel (*79*) using default parameters.

Docking was performed using AutoDock Vina with Hydrogens added to both ligand and receptor (*80*, *81*). To perform blind docking over the whole surface of each protein, a grid box perfectly framing each protein was generated with a custom python script and output as an AutoDock Vina config file. The exhaustiveness was scaled to the size of the search area (but was equivalent to 24 per 30x30x30 grid) and the number of poses was set to 30. Briefly, the 30 poses were re-scored using the RF-Score-Vs scoring function (*82*) but were found to have good agreement with the original AutoDock Vina scores. Using ChimeraX, the top 10 poses (based on original scores) for each receptor-ligand combination were then iterated through, and all residues within 5 Ångstroms of the ligand were output. The number of times each residue occurred was counted and added to a reference table like (described above).

#### Generation of expression plasmids

For plasmids expressing wild-type orthologues of Octβ2: The CDS from *Anopheles gambiae* was amplified using forward primer ttgtacaaaaaagcaggctgccaccATGATGAATCCTTCCAATGACGAC and reverse primer caccatcgaacctgatccAAGGGACTCTCCGATTTCAGAG. The CDS from *Apis mellifera* was amplified from cDNA extracted from honeybee heads (a kind gift from Mark J F Brown, Royal Holloway University) using forward primer ttgtacaaaaaagcaggctgccaccATGACGACGATCGTGACGAG and reverse primer caccatcgaacctgatccGAGGCTGCTACCGTACTCG. The CDS from *Varroa destructor* was synthesised with codon optimisation for mammalian expression (GENEWIZ) and amplified using forward primer ttgtacaaaaaagcaggctgccaccATGTCGGTAGAAGCGGGG and reverse primer caccatcgaacctgatccAGGATCATTTCTACAGGACGGTT. Each CDS was amplified without a stop codon and overhangs on each side complementary to plasmid backbone and mScarlet, to be used for Gibson assembly. The reverse primer also provided a GSGS linker between Octβ2 and mScarlet. The mScarlet CDS, including stop codon, was amplified using GTCCCTTggatcaggttcgatggtgagcaagggcgag (*An. gambiae*), CAGCCTCggatcaggttcgatggtgagcaagggcgag (*Ap. mellifera*) or CGACCCCggatcaggttcgatggtgagcaagggcgag (*V. destructor*) as a forward primer and cacactggactagtggatccgagctgtttacttgtacagctcgtccatg as a reverse primer. These two fragments were used in a three fragment Gibson assembly with a backbone digested from pVB210525 (XhoI, SacI), containing an IRES-Puromycin resistance cassette, a CAG promotor, an Ampicillin selective marker, and Tol2 Inverted Terminal Repeats.

For the point mutant constructs, the same approach as above was used, but each Octβ2 CDS was split into two fragments at the mutation site. The following reverse primers were used in conjunction with the original *An. gambiae* Octβ2 forward primer (above):

GAAAAGTAGACcatCAAAGAGTTCCACACATCACACA (D212M), TAACAATGAATtcGCAAGAGTCGGGGTTATCAAGA (L290E),

CTATGTCTGGGcaAGGACATCTATCGCACAGACTTG (N432C). The following forward primers were used in conjunction with the original *An. gambiae* Octβ2 reverse primer (above): GGAACTCTTTGatgGTCTACTTTTCCACTGCTAGCATC (D212M), CCCGACTCTTGCgaATTCATTGTTAACAAGCCTTATGCC (L290E), GATAGATGTCCTtgCCCAGACATAGTAGTAGCTTTAGTGT (N432C).

For the G-Flamp2 expression plasmid, we used a similar approach but performed a three-fragment assembly of G-Flamp2 CDS, IRES-NeoR (Neomycin/G418 resistance cassette) and a backbone digested from pVB210525 (XhoI, FseI), containing a CAG promotor (to drive G-Flamp2 expression), an Ampicillin selective marker and Tol2 Inverted Terminal Repeats. G-Flamp2 was amplified from pCAG-G-Flamp2 (AddGene #192782) with forward primer tgtacaaaaaagcaggctgccacccatgggctcccaccacca and reverse primer aaagctgggtctacctcattaggctgaggcagctg. The IRES-NeoR cassette was amplified from pVB180509 with forward primer agcctaatgaggtagacccagctttcttgtaca and reverse primer tatcatgtctgctcgaagcggccgggttgtcagaagaactcgtcaagaag. Restriction enzymes were purchased from NEB, NEBuilder HiFi DNA Assembly mix (NEB) was used for Gibson assembly and NEB Stable Competent *E. coli* (High Efficiency) cells were used for transformation. Plasmid DNA was prepared for transfection using Endotoxin-free maxiprep kits (Zymo).

#### Cell expression

HeLa cells were co-transfected with Tol2 transposase helper plasmid and G-Flamp2 expression plasmid using Lipofectamine 2000 (ThermoFisher) and Optimem (Gibco). Integrants were then selected with G418 (Sigma). This cell line was then separately co-transfected with transposase and each respective Octβ2 expression vector, each time selecting with Puromycin. Cells were cultured in high-glucose DMEM (Gibco) supplemented with 10% FBS (Gibco) at 37°C with 5% CO_2_. The day before an experiment, cells were seeded into black-walled optical-bottomed 96-well plates at approximately 10,000 cells per well. On the day of experiments, culture medium was exchanged for DMEM with 0.5% FBS and cells were serum starved for three hours. Amitraz was initially dissolved in DMSO, and octopamine in water – both were dissolved at 50 mM and 8x 10-fold serial dilutions were made in their respective solvent. These were then diluted to a 2x concentration in DMEM with 0.5% FBS. Wells were imaged with auto-focus using an automated stage on a Zeiss 880 Inverted Confocal. An image of each well was taken before the addition of drugs. Drugs were then added to 1x using a multichannel pipette, and wells were imaged again after ≥90 sec. Each well was imaged with a 10x plan apochromatic objective, laser levels and gain were kept constant for each given cell line.

#### Image analysis to determine cAMP responses

To identify cells and quantify their G-Flamp2 expression before and after drug treatment, we developed an image analysis pipeline in Python using OpenCV. This tool, named FLUOREST (FLUOrescent RESponse Timecourse analyser), is available on GitHub (https://github.com/mosquitome/fluorest). Briefly, it iterates through each well, importing a CZI file with “pre” and “post” images in red and green channels, and annotates wells based on user-provided metadata. Registration of pre and post images ensures the correct assignment of pixels to each cell across timepoints. Cells are then identified by performing contour detection on a thresholded image, with contours filtered by size. A summary showing images post-registration with cell outlines is produced for each well, allowing user verification. The output of FLUOREST is red and green fluorescence values (mean, median, sum and max), pre and post, for each cell.

Over 20,000 cells were analysed in total, with an average of 149.8 ± 95.9 (mean ± standard deviation) cells per treatment. Experiments were replicated at least three times. Cells with saturated pixels (i.e. a maximum fluorescence of 4095) at any timepoint were filtered. Median pre-drug green fluorescence was subtracted from median post-drug green fluorescence to give the change in fluorescence of each cell (ΔF). Four-parameter logistic models were then fit to data. Normalised fluorescence was calculated as

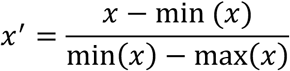

where min and max values correspond to the minima and maxima of the four-parameter logistic fit.

#### StaSsScal analysis

For LDV and electrophysiology experiments, statistical tests for normality (Shapiro-Wilk test with a significance level *of p < 0.05*) were used to test if data were normally distributed. These were generally found to be normally distributed; thus, mean and standard deviation values are reported throughout the manuscript. Data were analysed using Wilcoxon rank-sum tests or independent samples t-test depending on the data distribution.

Phonotaxis data were analysed using a binomial generalized linear model (GLM) using genotype and frequency as predictor variables. Insemination and eggs phenotype related data were analysed using binomial generalized linear model (GLM) including genotype as predictor variable.

Survival data were analysed using the Mantel–Cox test (log-rank test), which compares survival distributions across groups using a Chi-squared statistic.

Statistical tests were performed in R 4.2.2 or Python 3.9. Biorender, Adobe Illustrator and InkScape was used to generate explanatory graphics.

## Acknowledgments

The authors would like to thank Mark JF Brown for kindly providing *Apis mellifera* samples, Anwen Bullen for microscopy support and Lubna Maharelly for laboratory assistance. MA is part of the CSIC’s Global Health Plaform (PTI Salud Global).

## Funding

UK Research and Innovation under the Future Leaders Fellowship scheme grant MR/S015493/1 and MR/Y011732/1 (MA)

UCL Global Challenge Research Fund grant (MA) UCL Engagement Fund (MA)

Wellcome ISSF grant 204841/Z/16/Z (MA)

State Research Agency ATRAE Programme grant ATR2023-145654 (MA)

Plan Generación de Conocimiento 2024 PID2023-146360OA-I00 (MA)

UCL Research Culture Award (DAE, ST)

Marie Skłodowska Curie Actions Global Postdoctoral Fellowship, underwritten by the UKRI EP/Z001994/1: HORIZON-MSCA-2023-PF-01-01 (DAE)

Biotechnology and Biological Sciences Research Council BBSRC, BB/V007866/ (JTA) The Human Frontier Science Program grant RGP0033/2021 (JTA)

Tokai Pathways to Global Excellence (T-GEx), MEXT Strategic Professional Development Program for Young Researchers grant 0121an0002 (MPS)

This publication was also supported by the project Research Infrastructures for the control of vector-borne diseases (Infravec2), which has received funding from the European Union’s Horizon 2020 research and innovation program under grant agreement No 731060.

## Author contributions

Conceptualization: DAE, SM, RS, JTA and MA; Methodology, DAE, ST, JB, MPS, JTA and MA; Investigation, DAE, ST, JB, ED, ST, YYJX, MP, EF, DTD, TS, ML, GI, MP, ML, DT, WN; Funding Acquisition, DAE, ST, JTA and MA; Supervision, SM, RS, JTA and MA Writing—original draft: DAE. MA

## Data and materials availability

Protein sequences used in this study were retrieved from Uniprot, including Octβ2 orthologues from *An. gambiae* (F5HKI1), *Ap. mellifera* (A0A7M7GMY8) and *V. destructor* (A0A7M7KEY3). All other source data supporting the findings of this study are included in the manuscript or in the Supplementary Information. Custom code is available on GitHub at (https://github.com/mosquitome/buzz-feeder).

## References

1. T. malERA R. C. P. on I. and D. Resistance, malERA: An updated research agenda for insecticide and drug resistance in malaria elimination and eradication. PLOS Medicine 14, e1002450 (2017).

2. World Health Organization, UNICEF, Global vector control response 2017-2030. (2017).

3. World malaria report 2021. Geneva: World Health Organization; 2021. Licence: CC BY-NC-SA 3.0 IGO.

4. R. S. Lees, B. Knols, R. Bellini, M. Q. Benedict, A. Bheecarry, H. C. Bossin, D. D. Chadee, J. Charlwood, R. K. Dabiré, L. Djogbenou, A. Egyir-Yawson, R. Gato, L. C. Gouagna, M. M. Hassan, S. A. Khan, L. L. Koekemoer, G. Lemperiere, N. C. Manoukis, R. Mozuraitis, R. J. Pitts, F. Simard, J. R. L. Gilles, Review: Improving our knowledge of male mosquito biology in relation to genetic control programmes. Acta Tropica 132, S2–S11 (2014).

5. A. Diabate, F. Tripet, Targeting male mosquito mating behaviour for malaria control. Parasites & Vectors 8, 347 (2015).

6. P. Barreaux, A. M. G. Barreaux, E. D. Sternberg, E. Suh, J. L. Waite, S. A. Whitehead, M. B. Thomas, Priorities for Broadening the Malaria Vector Control Tool Kit. Trends in Parasitology 33, 763–774 (2017).

7. M. V. Murgia, S. Sharan, J. Kaur, W. Austin, L. Hagen, L. Wu, L. Chen, J. A. Scott, D. P. Flaherty, M. E. Scharf, V. J. Watts, C. A. Hill, High-content phenotypic screening identifies novel chemistries that disrupt mosquito activity and development. Pestic Biochem Physiol 182, 105037 (2022).

8. L. Bi, A. F. Oliveros-Diaz, M. V. Murgia, J. Kaur, W. Austin, L. Wu, L. Chen, A. D. Gondhalekar, M. E. Scharf, C. A. Hill, High-content phenotypic screening reveals natural product bioactives that disrupt mosquito activity, development and serotonergic signaling. Pesticide Biochemistry and Physiology 213, 106498 (2025).

9. A. Hammond, R. Galizi, K. Kyrou, A. Simoni, C. Siniscalchi, D. Katsanos, M. Gribble, D. Baker, E. Marois, S. Russell, A. Burt, N. Windbichler, A. Crisanti, T. Nolan, A CRISPR-Cas9 Gene Drive System Targeting Female Reproduction in the Malaria Mosquito vector Anopheles gambiae. Nat Biotechnol 34, 78–83 (2016).

10. K. Kyrou, A. M. Hammond, R. Galizi, N. Kranjc, A. Burt, A. K. Beaghton, T. Nolan, A. Crisanti, A CRISPR–Cas9 gene drive targeting doublesex causes complete population suppression in caged Anopheles gambiae mosquitoes. Nature biotechnology (2018).

11. I. Tolosana, K. Willis, M. Gribble, L. Phillimore, A. Burt, T. Nolan, A. Crisanti, F. Bernardini, A Y chromosome-linked genome editor for efficient population suppression in the malaria vector Anopheles gambiae. Nat Commun 16, 206 (2025).

12. A. N. Clements, The biology of mosquitoes. Sensory reception and behaviour. 1999. Wallingford: CABI Publishing Google Scholar.

13. A. Diabaté, A. S. Yaro, A. Dao, M. Diallo, D. L. Huestis, T. Lehmann, Spatial distribution and male mating success of Anopheles gambiae swarms. BMC Evolutionary Biology 11, 184 (2011).

14. L. J. Cator, B. J. Arthur, L. C. Harrington, R. R. Hoy, Harmonic convergence in the love songs of the dengue vector mosquito. Science 323, 1077–9 (2009).

15. C. Pennetier, B. Warren, K. R. Dabiré, I. J. Russell, G. Gibson, “Singing on the wing” as a mechanism for species recognition in the malarial mosquito Anopheles gambiae. Current Biology 20, 131–136 (2010).

16. B. Warren, G. Gibson, I. J. Russell, Sex Recognition through midflight mating duets in Culex mosquitoes is mediated by acoustic distortion. Current Biology 19, 485–491 (2009).

17. J. Somers, M. Georgiades, M. P. Su, J. Bagi, M. Andrés, A. Alampounti, G. Mills, W. Ntabaliba, S. J. Moore, R. Spaccapelo, J. T. Albert, Hitting the right note at the right time: Circadian control of audibility in Anopheles mosquito mating swarms is mediated by flight tones. Sci Adv 8, eabl4844 (2022).

18. M. Andrés, M. P. Su, J. Albert, L. J. Cator, Buzzkill: targeting the mosquito auditory system. Current Opinion in Insect Science 40, 11–17 (2020).

19. Y. M. Loh, M. P. Su, D. A. Ellis, M. Andrés, The auditory efferent system in mosquitoes. Frontiers in Cell and Developmental Biology 11 (2023).

20. E. A. Freeman, D. A. Ellis, J. Bagi, S. Tytheridge, M. Andrés, Perspectives on the manipulation of mosquito hearing. Current Opinion in Insect Science 66, 101271 (2024).

21. Y. Wang, D. Thakur, E. Duge, C. Murphy, I. Girling, N. A. DeBeaubien, J. Chen, B. H. Nguyen, A. S. Gurav, C. Montell, Deafness due to loss of a TRPV channel eliminates mating behavior in Aedes aegypti males. Proceedings of the National Academy of Sciences 121, e2404324121 (2024).

22. M. Georgiades, A. Alampounti, J. Somers, M. P. Su, D. A. Ellis, J. Bagi, D. Terrazas-Duque, S. Tytheridge, W. Ntabaliba, S. Moore, J. T. Albert, M. Andrés, Hearing of malaria mosquitoes is modulated by a beta-adrenergic-like octopamine receptor which serves as insecticide target. Nat Commun 14, 4338 (2023).

23. T. Roeder, Tyramine and octopamine: ruling behavior and metabolism. Annu Rev Entomol 50, 447–477 (2005).

24. P. V. Pietrantonio, C. Xiong, R. J. Nachman, Y. Shen, G protein-coupled receptors in arthropod vectors: omics and pharmacological approaches to elucidate ligand-receptor interactions and novel organismal functions. Current opinion in insect science (2018).

25. N. Audsley, R. E. Down, G protein coupled receptors as targets for next generation pesticides. Insect Biochemistry and Molecular Biology 67, 27–37 (2015).

26. C. A. Hill, S. Sharan, V. J. Watts, Genomics, GPCRs and new targets for the control of insect pests and vectors. Current Opinion in Insect Science 30, 99–106 (2018).

27. T. Kita, T. Hayashi, T. Ohtani, H. Takao, H. Takasu, G. Liu, H. Ohta, F. Ozoe, Y. Ozoe, Amitraz and its metabolite differentially activate α- and β-adrenergic-like octopamine receptors. Pest Management Science 73, 984–990 (2017).

28. P. Vudriko, J. Okwee-Acai, J. Byaruhanga, D. S. Tayebwa, S. G. Okech, R. Tweyongyere, E. M. Wampande, A. R. A. Okurut, K. Mugabi, J. B. Muhindo, J. L. Nakavuma, R. Umemiya-Shirafuji, X. Xuan, H. Suzuki, Chemical tick control practices in southwestern and northwestern Uganda. Ticks and Tick-borne Diseases 9, 945–955 (2018).

29. N. Pugliese, E. Circella, G. Cocciolo, A. Giangaspero, D. H. Tomic, T. S. Kika, A. Caroli, A. Camarda, Efficacy of λ-cyhalothrin, amitraz, and phoxim against the poultry red mite Dermanyssus gallinae De Geer, 1778 (Mesostigmata: Dermanyssidae): an eight-year survey. Avian Pathology 48, S35–S43 (2019).

30. L. Guo, X. Fan, X. Qiao, C. Montell, J. Huang, An octopamine receptor confers selective toxicity of amitraz on honeybees and Varroa mites. eLife 10, e68268 (2021).

31. M. a. I. Ahmed, F. Matsumura, Synergistic actions of formamidine insecticides on the activity of pyrethroids and neonicotinoids against Aedes aegypti (Diptera: Culicidae). J Med Entomol 49, 1405–1410 (2012).

32. M. A. I. Ahmed, C. F. A. Vogel, Synergistic action of octopamine receptor agonists on the activity of selected novel insecticides for control of dengue vector Aedes aegypti (Diptera: Culicidae) mosquito. Pesticide Biochemistry and Physiology 120, 51–56 (2015).

33. P. M. V. Simões, R. A. Ingham, G. Gibson, I. J. Russell, A role for acoustic distortion in novel rapid frequency modulation behaviour in free-flying male mosquitoes. The Journal of Experimental Biology 219, 2039–2047 (2016).

34. S. Fuchs, E. Rende, A. Crisanti, T. Nolan, Disruption of aminergic signalling reveals novel compounds with distinct inhibitory effects on mosquito reproduction, locomotor function and survival. Scientific Reports 4, srep05526 (2014).

35. J. Lim, P. R. Sabandal, A. Fernandez, J. M. Sabandal, H.-G. Lee, P. Evans, K.-A. Han, The Octopamine Receptor Octβ2R Regulates Ovulation in Drosophila melanogaster. PLOS ONE 9, e104441 (2014).

36. Y. Li, C. Fink, S. El-Kholy, T. Roeder, THE OCTOPAMINE RECEPTOR octß2R IS ESSENTIAL FOR OVULATION AND FERTILIZATION IN THE FRUIT FLY Drosophila melanogaster. Archives of Insect Biochemistry and Physiology 88, 168–178 (2015).

37. M. A. White, D. S. Chen, M. F. Wolfner, She’s got nerve: roles of octopamine in insect female reproduction. J Neurogenet 35, 132–153 (2021).

38. H. Ohta, Y. Ozoe, “Chapter Two - Molecular Signalling, Pharmacology, and Physiology of Octopamine and Tyramine Receptors as Potential Insect Pest Control Targets” in Advances in Insect Physiology, E. Cohen, Ed. (Academic Press, 2014; https://www.sciencedirect.com/science/article/pii/B9780124170100000021)vol. 46 of *Target Receptors in the Control of Insect Pests: Part II*, pp. 73–166.

39. Guidelines for laboratory and field testing of long-lasting insecticidal nets. https://www.who.int/publications/i/item/9789241505277.

40. L. Facchinelli, L. Valerio, R. S. Lees, C. F. Oliva, T. Persampieri, C. M. Collins, A. Crisanti, R. Spaccapelo, M. Q. Benedict, Stimulating Anopheles gambiae swarms in the laboratory: application for behavioural and fitness studies. Malaria Journal 14, 271 (2015).

41. K. W. Kastner, D. A. Shoue, G. L. Estiu, J. Wolford, M. F. Fuerst, L. D. Markley, J. A. Izaguirre, M. A. McDowell, Characterization of the Anopheles gambiae octopamine receptor and discovery of potential agonists and antagonists using a combined computational-experimental approach. Malaria Journal 13, 434 (2014).

42. W. Liu, C. Liu, P.-G. Ren, J. Chu, L. Wang, An Improved Genetically Encoded Fluorescent cAMP Indicator for Sensitive cAMP Imaging and Fast Drug Screening. Front. Pharmacol. 13 (2022).

43. J. Huang, T. Hamasaki, F. Ozoe, H. Ohta, K. Enomoto, H. Kataoka, Y. Sawa, A. Hirota, Y. Ozoe, Identification of Critical Structural Determinants Responsible for Octopamine Binding to the α-Adrenergic-like Bombyx mori Octopamine Receptor. Biochemistry 46, 5896–5903 (2007).

44. L. M. Roth, A study of mosquito behavior. An experimental laboratory study of the sexual behavior of Aedes aegypti (Linnaeus). The American Midland Naturalist 40, 265–352 (1948).

45. P. Belton, Attraction of male mosquitoes to sound. J. Am. Mosq. Control 10, 297–301 (1994).

46. M. P. Su, M. Andrés, N. Boyd-Gibbins, J. Somers, J. T. Albert, Sex and species specific hearing mechanisms in mosquito flagellar ears. Nat Commun. 9, 3911 (2018).

47. K. S. Boo, Antennal sensory receptors of the male mosquito, Anopheles stephensi. Zeitschrift für Parasitenkunde 61, 249–264 (1980).

48. K. S. Boo, A. G. Richards, Fine structure of the scolopidia in the Johnston’s organ of male Aedes aegypti (L.)(Diptera: Culicidae). International Journal of Insect Morphology and Embryology 4, 549–566 (1975).

49. K. S. Boo, A. G. Richards, Fine structure of scolopidia in Johnston’s organ of female Aedes aegypti compared with that of the male. Journal of Insect Physiology 21, 1129–1139 (1975).

50. M. Andrés, M. Seifert, C. Spalthoff, B. Warren, L. Weiss, D. Giraldo, M. Winkler, S. Pauls, M. C. Göpfert, Auditory Efferent System Modulates Mosquito Hearing. Curr Biol 26, 2028– 2036 (2016).

51. D. D. Vorontsov, D. N. Lapshin, Effect of Octopamine on the Frequency Tuning of the Auditory System in Culex Pipiens Pipiens Mosquito (Diptera, Culicidae). Neurosci Behav Physi 54, 319–328 (2024).

52. Y. Y. J. Xu, Y. M. Loh, T.-T. Lee, W.-T. Chen, W. Loh, T. S. Ohashi, D. F. Eberl, M. Andrés, M. P. Su, A. Kamikouchi, cAMP-related second messenger pathways modulate hearing function in *Aedes aegypti* mosquitoes. iScience, 113202 (2025).

53. A. Widmer, U. Höger, S. Meisner, A. S. French, P. H. Torkkeli, Spider Peripheral Mechanosensory Neurons Are Directly Innervated and Modulated by Octopaminergic Efferents. J. Neurosci. 25, 1588–1598 (2005).

54. N. e. Gruntenko, E. k. Karpova, A. a. Alekseev, N. a. Chentsova, E. v. Bogomolova, M. Bownes, I. Yu. Rauschenbach, Effects of octopamine on reproduction, juvenile hormone metabolism, dopamine, and 20-hydroxyecdysone contents in Drosophila. Archives of Insect Biochemistry and Physiology 65, 85–94 (2007).

55. M. Salomon, O. Malka, R. K. V. Meer, A. Hefetz, The role of tyramine and octopamine in the regulation of reproduction in queenless worker honeybees. Naturwissenschaften 99, 123– 131 (2012).

56. J. Leyria, I. Orchard, A. B. Lange, Octopamine is required for successful reproduction in the classical insect model, Rhodnius prolixus. PLOS ONE 19, e0306611 (2024).

57. D. Liu, X. Zhang, F. Chiqin, I. Nyamwasa, Y. Cao, J. Yin, S. Zhang, H. Feng, K. Li, Octopamine modulates insect mating and Oviposition. J Chem Ecol 48, 628–640 (2022).

58. H.-G. Lee, C.-S. Seong, Y.-C. Kim, R. L. Davis, K.-A. Han, Octopamine receptor OAMB is required for ovulation in Drosophila melanogaster. Developmental Biology 264, 179–190 (2003).

59. E. W. Rohrbach, E. M. Knapp, S. A. Deshpande, D. E. Krantz, Expression and potential regulatory functions of Drosophila octopamine receptors in the female reproductive tract. G3 (Bethesda) 14, jkae012 (2024).

60. S. A. Deshpande, E. W. Rohrbach, J. D. Asuncion, J. Harrigan, A. Eamani, E. H. Schlingmann, D. J. Suto, P.-T. Lee, F. E. Schweizer, H. J. Bellen, D. E. Krantz, Regulation of Drosophila oviduct muscle contractility by octopamine. iScience 25, 104697 (2022).

61. F. W. Avila, M. C. Bloch Qazi, C. D. Rubinstein, M. F. Wolfner, A requirement for the neuromodulators octopamine and tyramine in Drosophila melanogaster female sperm storage. Proceedings of the National Academy of Sciences 109, 4562–4567 (2012).

62. C. A. Middleton, U. Nongthomba, K. Parry, S. T. Sweeney, J. C. Sparrow, C. J. Elliott, Neuromuscular organization and aminergic modulation of contractions in the Drosophila ovary. BMC Biology 4, 17 (2006).

63. E. M. Walsh, M. A. Janowiecki, K. Zhu, N. H. Ing, E. L. Vargo, J. Rangel, Elevated Mating Frequency in Honey Bee (Hymenoptera: Apidae) Queens Exposed to the Miticide Amitraz During Development. Annals of the Entomological Society of America 114, 620–626 (2021).

64. R. Baeshen, N. E. Ekechukwu, M. Toure, D. Paton, M. Coulibaly, S. F. Traoré, F. Tripet, Differential effects of inbreeding and selection on male reproductive phenotype associated with the colonization and laboratory maintenance of Anopheles gambiae. Malar J 13, 19 (2014).

65. Á. Lehane, C. Parker-Crockett, E. Norris, S. S. Wheeler, L. C. Harrington, Measuring insecticide resistance in a vacuum: exploring next steps to link resistance data with mosquito control efficacy. Journal of Medical Entomology 61, 584–594 (2024).

66. A. S. Probst, D. G. Paton, F. Appetecchia, S. Bopp, K. L. Adams, T. A. Rinvee, S. Pou, R. Winter, E. W. Du, S. Yahiya, C. Vidoudez, N. Singh, J. Rodrigues, P. Castañeda-Casado, C. Tammaro, D. Chen, K. P. Godinez-Macias, J. L. Jaramillo, G. Poce, M. J. Rubal, A. Nilsen, E. A. Winzeler, J. Baum, J. N. Burrows, M. K. Riscoe, D. F. Wirth, F. Catteruccia, In vivo screen of Plasmodium targets for mosquito-based malaria control. Nature 643, 785–793 (2025).

67. R. A. Baines, M. Bate, Electrophysiological Development of Central Neurons in the Drosophila Embryo. J. Neurosci. 18, 4673–4683 (1998).

68. A. Buslaev, V. I. Iglovikov, E. Khvedchenya, A. Parinov, M. Druzhinin, A. A. Kalinin, Albumentations: Fast and Flexible Image Augmentations. Information 11, 125 (2020).

69. J. T. Albert, B. Nadrowski, M. C. Göpfert, Mechanical Signatures of Transducer Gating in the Drosophila Ear. Current Biology 17, 1000–1006 (2007).

70. A. Cavagna, I. Giardina, M. A. Gucciardino, G. Iacomelli, M. Lombardi, S. Melillo, G. Monacchia, L. Parisi, M. J. Peirce, R. Spaccapelo, Characterization of lab-based swarms of Anopheles gambiae mosquitoes using 3D-video tracking. Sci Rep 13, 8745 (2023).

71. H. M. Ferguson, K. R. Ng’habi, T. Walder, D. Kadungula, S. J. Moore, I. Lyimo, T. L. Russell, H. Urassa, H. Mshinda, G. F. Killeen, B. G. Knols, Establishment of a large semi-field system for experimental study of African malaria vector ecology and control in Tanzania. Malaria Journal 7, 158 (2008).

72. J. D. Charlwood, M. D. R. Jones, Mating in the mosquito, Anopheles gambiae s.l. Physiological Entomology 5, 315–320 (1980).

73. M. Q. Benedict, Methods in Anopheles research. Malaria research and reference reagent resource center (MR4) (2007).

74. C. F. Oliva, M. Q. Benedict, G. Lempérière, J. Gilles, Laboratory selection for an accelerated mosquito sexual development rate. Malar J 10, 135 (2011).

75. E. W. Kaindoa, H. S. Ngowo, A. J. Limwagu, M. Tchouakui, E. Hape, S. Abbasi, J. Kihonda, A. S. Mmbando, R. M. Njalambaha, G. Mkandawile, H. Bwanary, M. Coetzee, F. O. Okumu, Swarms of the malaria vector Anopheles funestus in Tanzania. Malaria Journal 18, 29 (2019).

76. J. Jumper, R. Evans, A. Pritzel, T. Green, M. Figurnov, O. Ronneberger, K. Tunyasuvunakool, R. Bates, A. Žídek, A. Potapenko, A. Bridgland, C. Meyer, S. A. A. Kohl, A. J. Ballard, A. Cowie, B. Romera-Paredes, S. Nikolov, R. Jain, J. Adler, T. Back, S. Petersen, D. Reiman, E. Clancy, M. Zielinski, M. Steinegger, M. Pacholska, T. Berghammer, S. Bodenstein, D. Silver, O. Vinyals, A. W. Senior, K. Kavukcuoglu, P. Kohli, D. Hassabis, Highly accurate protein structure prediction with AlphaFold. Nature 596, 583–589 (2021).

77. M. Varadi, S. Anyango, M. Deshpande, S. Nair, C. Natassia, G. Yordanova, D. Yuan, O. Stroe, G. Wood, A. Laydon, A. Žídek, T. Green, K. Tunyasuvunakool, S. Petersen, J. Jumper, E. Clancy, R. Green, A. Vora, M. Lutfi, M. Figurnov, A. Cowie, N. Hobbs, P. Kohli, G. Kleywegt, E. Birney, D. Hassabis, S. Velankar, AlphaFold Protein Structure Database: massively expanding the structural coverage of protein-sequence space with high-accuracy models. Nucleic Acids Research 50, D439–D444 (2022).

78. J. J. Irwin, K. G. Tang, J. Young, C. Dandarchuluun, B. R. Wong, M. Khurelbaatar, Y. S. Moroz, J. Mayfield, R. A. Sayle, ZINC20—A Free Ultralarge-Scale Chemical Database for Ligand Discovery. J. Chem. Inf. Model. 60, 6065–6073 (2020).

79. N. M. O’Boyle, M. Banck, C. A. James, C. Morley, T. Vandermeersch, G. R. Hutchison, Open Babel: An open chemical toolbox. Journal of Cheminformatics 3, 33 (2011).

80. J. Eberhardt, D. Santos-Martins, A. F. Tillack, S. Forli, AutoDock Vina 1.2.0: New Docking Methods, Expanded Force Field, and Python Bindings. J. Chem. Inf. Model. 61, 3891–3898 (2021).

81. O. Trott, A. J. Olson, AutoDock Vina: Improving the speed and accuracy of docking with a new scoring function, efficient optimization, and multithreading. Journal of Computational Chemistry 31, 455–461 (2010).

82. M. Wójcikowski, P. J. Ballester, P. Siedlecki, Performance of machine-learning scoring functions in structure-based virtual screening. Sci Rep 7, 46710 (2017).

83. E. C. Meng, T. D. Goddard, E. F. Pettersen, G. S. Couch, Z. J. Pearson, J. H. Morris, T. E. Ferrin, UCSF ChimeraX: Tools for structure building and analysis. Protein Science 32, e4792 (2023).

