## Supplementary material for "Controlling malaria mosquito reproduction via the octopamine beta2 receptor"

Fig. S1.

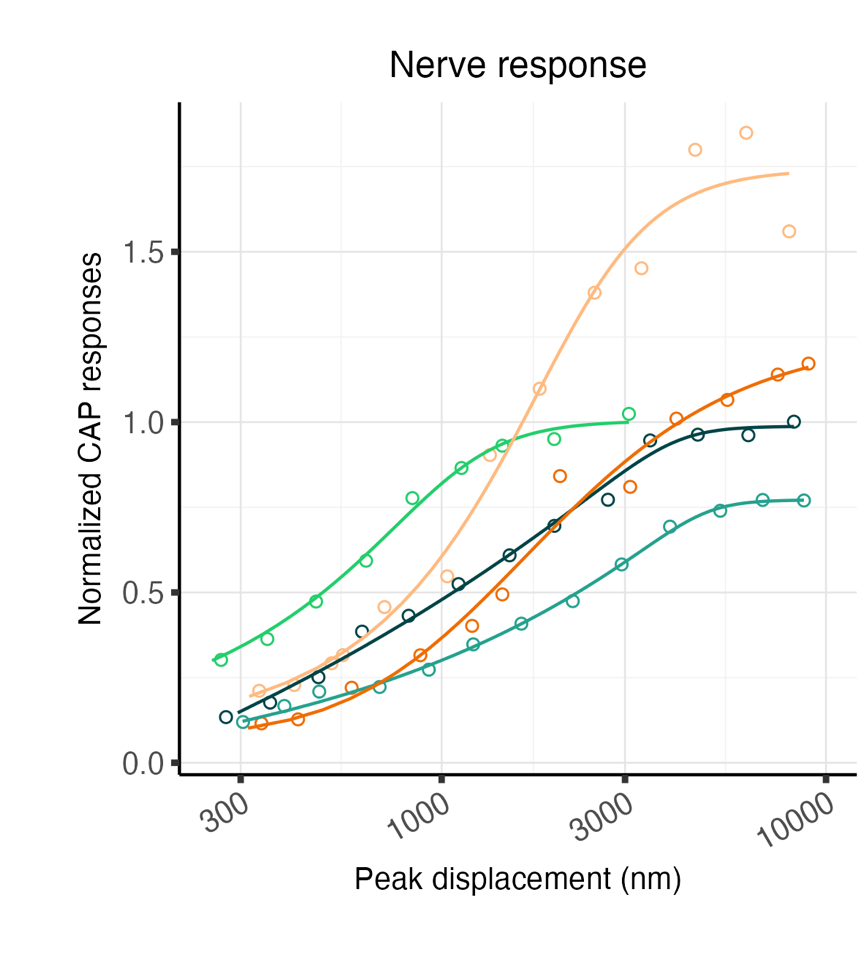

**Supplementary Figure 1: Raw antennal nerve CAP responses in response to flagellar displacements (5 parameter log-logistic fits).** An increase in CAP amplitude relative to flagellar displacement values is also observed upon octopamine injections in wt individuals when comparing raw CAP voltage values (rather than normalized CAP values as in Fig.1h). Samples sizes= 8. CAP compound action potentials.

Fig. S2.

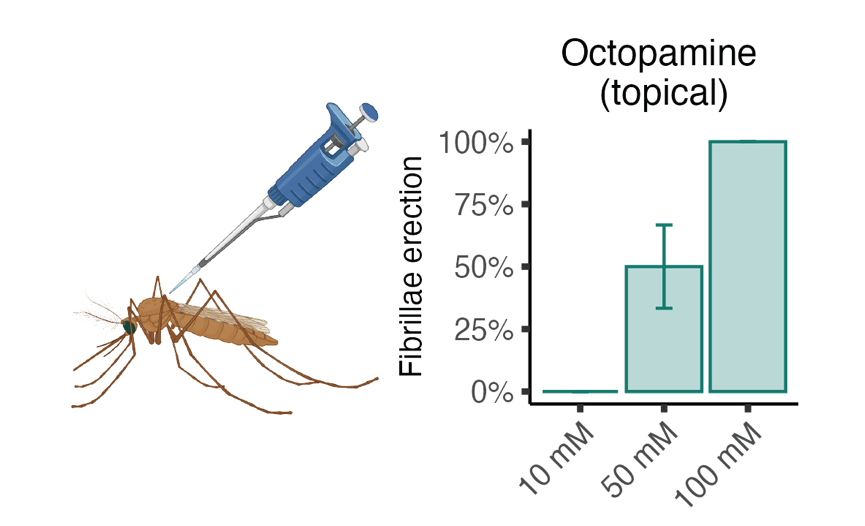

**Supplementary Figure 2: Topical exposure can also be used to administer compounds to mosquitoes, but their efficacy is lower than compound feeding or injection.** 100 mM octopamine dissolved in 80% DMSO were required to cause fibrillae erection in 100% male mosquitoes. Three replicates were conducted, 5 mosquitoes were tested in each replicate.

Fig. S3.

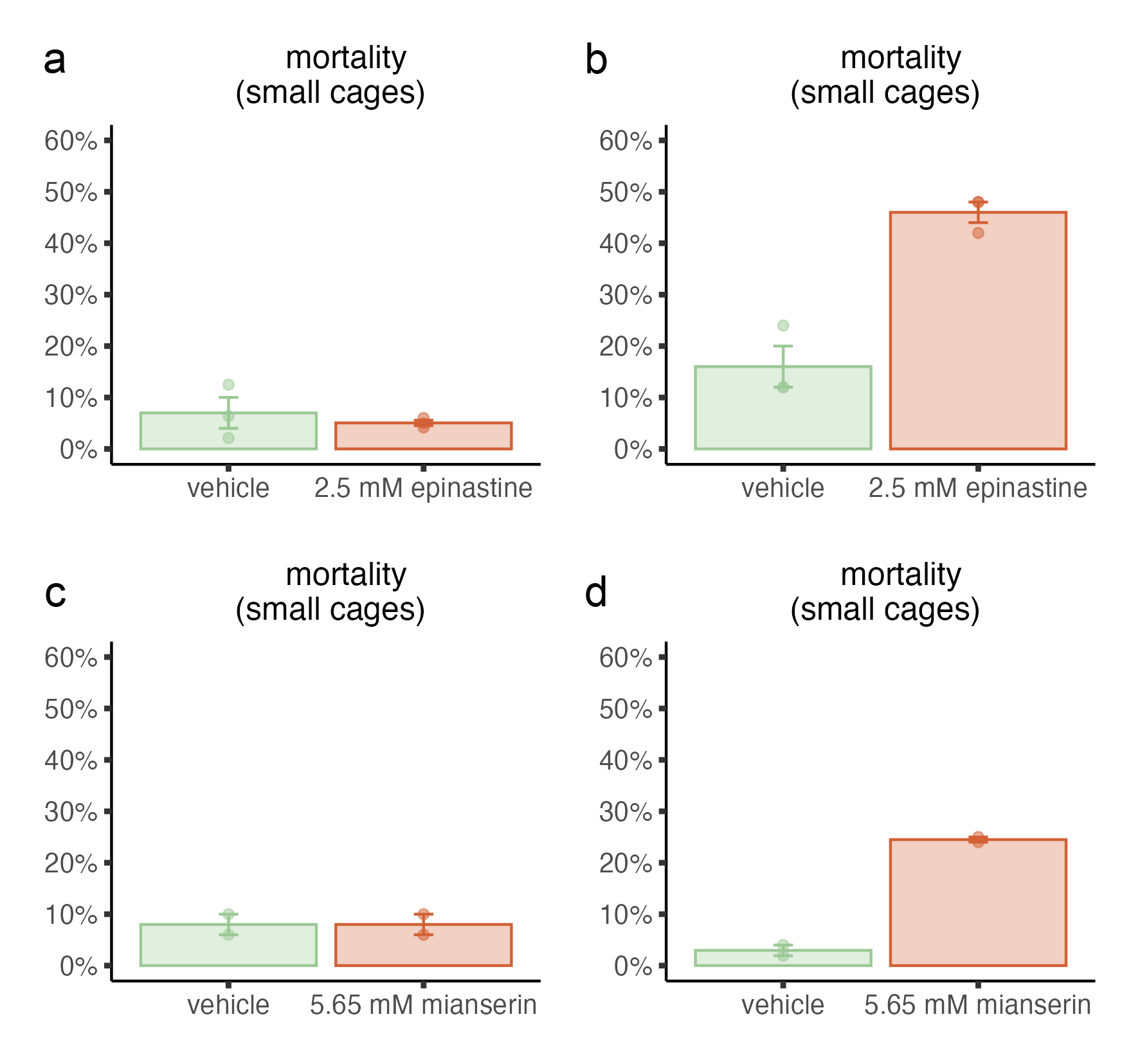

**Supplementary Figure 3: Feeding mosquitoes with beta-adrenergic-like octopamine receptor antagonists cause high mortality across males.** Male and female mosquitoes were fed either 5 mM mianserin or 2.5mM epinastine as in Fig. 5a. After 22 hours (one swarming period), male and female mortality was assessed. The mortality across males was very high upon feeding in either antagonist (**b,d**; epinastine: 46%; mianserin: 24%), making it difficult to interpret the mating behaviour assay results. Each dot represents the mean of a replicate using 50 females in the initial crosses, total of 3 (epinastine) and 2 (mianserin) replicates.

Table S1.

| Parameter | sex | Genotype | Condition | mean ± SD | p.adj (pairwise comparisons) |
| --- | --- | --- | --- | --- | --- |
| Mechanical tuning (Hz) | female | wt | baseline | 281.9  48.9 | Wilcoxon rank-sum test:  Baseline-control: 0.49;  Control-1mM OA: 0.47;  Baseline-1mM OA: 0.83 |
|  | female | wt | control | 285.6  59.6 |  |
|  | female | wt | 1mM OA | 305.0  112.1 |  |
|  | male | wt | baseline | 358.2  42.7 | Independent samples t-tests:  Baseline-control: 0.64;  Control-1mM OA: 0.014*;  Baseline-1mM OA: 0.014*. |
|  | male | wt | control | 366.5  51.8 |  |
|  | male | wt | 1mM OA | 533.5  123.5 |  |
|  | male | *AgOctβ2^-^* | baseline | 378.0  17.8 | Wilcoxon rank-sum test:  Baseline-1mM OA: 0.38 |
|  | male | *AgOctβ2^-^* | 1mM OA | 359.3  28.1 |  |
| Electrical tuning (Hz) | female | wt | baseline | 209.1  13.4 | Wilcoxon rank-sum test:  Baseline-control: 0.92;  Control-1mM OA: 0.47;  Baseline-1mM OA: 0.72 |
|  | female | wt | control | 237.2  68.432009 |  |
|  | female | wt | 1mM OA | 221.1  71.5 |  |
|  | male | wt | baseline | 189.9  23.5 | Independent samples t-tests:  Baseline-control: 0.04;  Control-1mM OA: 0.000002****;  Baseline-1mM OA: 0.000016**** |
|  | male | wt | control | 166.0  28.9 |  |
|  | male | wt | 1mM OA | 270.8500  26.7 |  |
|  | male | *AgOctβ2^-^* | baseline | 234.6  35.5 | Independent samples t-tests:  Baseline-1mM OA: 0.42 |
|  | male | *AgOctβ2^-^* | 1mM OA | 218.9  34.4 |  |

**Supplementary Table 1: Summary of auditory tuning frequencies in male and female, wild-type and *AgOctβ2^-^* An. gambiae, before and after octopamine injection.** OA octopamine.

Fig. S4 (separate file)

STL file containing feeding assay plate for 3D-printing.
